# A Nonlinear Dynamics Framework for Mutation-Selection Balance Reveals Dominance-Dependent Bistability in Auto- and Allotetraploids

**DOI:** 10.1101/2025.08.28.672890

**Authors:** Samuel R. Gibbon, Justin L. Conover, Michael S. Barker, Ryan N. Gutenkunst

**Author notes:** Department of Human Genetics, University of Chicago, Chicago, IL, 60637, USA.

## Abstract

Polyploidy and whole genome duplication (WGD) are widespread biological phenomena with substantial cellular, meiotic, and genetic effects. Despite their prevalence and significance across the tree of life, population genetics theory for polyploids is not well developed. The lack of theoretical models limits our understanding of polyploid evolution and restricts our ability to harness polyploidy for crop improvement amidst increasing environmental stress. To address this gap, we developed and analyzed deterministic models of mutation-selection balance for tetraploids under polysomic (autotetraploid) and disomic (allotetraploid) inheritance patterns and arbitrary dominance relationships. We also introduced a new mathematical framework based on ordinary differential equations and nonlinear dynamics for analyzing the models. We find that the autotetraploid and allotetraploid models generally exhibit negligible differences in mutation load and allele frequency at mutation-selection balance. Our model also reveals two bistable points of mutation-selection balance for dominant alleles with biased mutation rates over a wide range of selection coefficients in the tetraploid models compared to bistability in only a narrow range for diploids. In addition, for overdominant mutations, we find a distinct equilibrium structure with bistable equilibria in only the allotetraploid model. Finally, using discrete time simulations, we explore the temporal dynamics of allele frequency and fitness change and show that the presence of bistability and bifurcations fundamentally alters the temporal dynamics of allele frequency and load. Overall, this work develops foundational theoretical models that will facilitate the development of population genetic models and methodologies to study evolution in empirical tetraploid populations.

## 1 Introduction

Whole genome duplication (polyploidy) represents one of the most dramatic mutational events in evolution, instantaneously doubling chromosome number and fundamentally altering meiotic (Lloyd and Bomblies, 2016), cellular (Fernandes Gyorfy et al., 2021), and evolutionary processes (Otto, 2007; Van De Peer et al., 2017). This genomic restructuring poses an apparent paradox: while both theory and empirical population genomic analysis find evidence for the reduced efficacy of selection in polyploids, accumulation of deleterious mutations, and higher mutation load at mutation-selection balance (Otto and Whitton, 2000; Otto, 2007; Douglas et al., 2015; Monnahan et al., 2019; Vlček et al., 2025), polyploid lineages flourish across the tree of life, especially among flowering plants (Mayrose et al., 2011; Arrigo and Barker, 2012; Van De Peer et al., 2017; One Thousand Plant Transcriptomes Initiative, 2019; McKibben et al., 2024). This widespread success may be shaped by the extent to which polyploidy reshapes mutation-selection dynamics, particularly through the masking of recessive deleterious mutations which reduces mutation load in polyploids (Otto and Whitton, 2000). However, quantitative frameworks for predicting evolutionary equilibria in polyploids remain incomplete, especially for allopolyploids with disomic inheritance that comprise many polyploid species (Barker et al., 2016; Dunn and Sethuraman, 2024). Despite the broad implications of polyploidy for agriculture (Renny-Byfield and Wendel, 2014; Sattler et al., 2016; Trojak-Goluch et al., 2021), its role in adaptation to environmental stress (Van De Peer et al., 2021; Turcotte et al., 2024; Blake-Mahmud et al., 2025), and its emerging relevance to cancer biology (Bielski et al., 2018; Matsumoto et al., 2021), we lack unified theoretical models that can predict how the balance between mutation and selection varies with ploidy, inheritance patterns, and dominance — a gap that limits our ability to understand foundational evolutionary processes in polyploids.

Polyploids are broadly categorized as either autopolyploid or allopolyploid based on their underlying genetics, chromosome pairing behavior during meiosis, or whether the polyploid formed as a result of interspecific hybridization (although these categories are often a false dichotomy (Twyford et al., 2025)). This distinction between auto- and allopolyploids has substantially shaped both empirical research (Barker et al., 2016; Doyle and Sherman-Broyles, 2017) and theoretical modeling (Haldane, 1926; Wright, 1938; Hill, 1970). Here, we adopt the genetic and cytological definitions which group auto- and allopolyploids into those with polysomic and disomic inheritance, respectively (Ramsey and Schemske, 2002; Parisod et al., 2010). Polysomic inheritance occurs when all of the chromosomes can pair in meiosis at random (i.e., with no preferential pairing) so that all possible chromosome pairing combinations occur with equal probability. Alternatively, disomic inheritance occurs when there are clearly distinguishable pairs of chromosomes (i.e., subgenomes) that always pair together and have inheritance patterns similar to that of diploids. Thus, under disomic inheritance, some allelic combinations in the resulting gametes do not occur because alleles in the same subgenome cannot segregate into the same gamete (e.g., sites with fixed heterozygosity).

Although models for polyploids have advanced to account for gametic/panmictic disequilibrium (Rowe, 1982; Rowe and Hill, 1984), population structure (Ronfort et al., 1998), double reduction (Butruille and Boiteux, 2000; Huang et al., 2019), selective sweeps (Monnahan and Brandvain, 2020), stabilizing selection acting on quantitative traits (Clo, 2022), and multiple loci (Griswold and Williamson, 2017; Mostafaee and Griswold, 2019; Griswold and Asif, 2023), all of these advances assume polysomic inheritance patterns (i.e., autopolyploidy) and do not consider disomic inheritance (i.e., allopolyploidy). This may be because each subgenome or pair of homologs under disomic inheritance can, in principle, be treated as a diploid which connects allopolyploids to the wealth of population genetics theory and computational/bioinformatic methods developed for diploids. This treatment greatly simplifies bioinformatic, genomic, and inference analyses (e.g., read mapping and genotyping) (Paape et al., 2016, 2018; Blischak et al., 2023); however, this approach also obscures fundamental evolutionary processes, such as selection, that operate on the integrated phenotype arising from the combination of subgenomes and other features of polyploids such as subgenome dominance and homoeolog expression bias (Grover et al., 2012; Wang et al., 2022; Zhang et al., 2023) or masking effects (Vekemans et al., 2026). Additionally, treating the subgenomes of allopolyploids as independent diploids assumes *a priori* that no genetic variation is co-segregating in both subgenomes; however, this assumption is rarely tested in practice (Twyford et al., 2025). These assumptions are most likely violated in newly-formed polyploids with recently diverged diploid progenitors (the relevant degree of divergence depending on the population sizes of the progenitor diploid populations), in cases where there is ongoing gene flow between the diploid progenitors prior to polyploid formation, or in systems where homoeologous exchanges occur, such as *Brassica napus* (Wang et al., 2024) or *Arachis hypogaea* (Lamon et al., 2026). Despite being equally as or more common in the empirical literature (Soltis et al., 2016; Barker et al., 2016; Spoelhof et al., 2017), allopolyploids have received less theoretical attention compared to autotetraploids (but see Schneemann and Welch, 2026). As a result, key questions about the dynamics of mutation and natural selection in disomic polyploids remain unanswered. Understanding these dynamics is important not only for insight into the evolutionary trajectories of allopolyploids, but also for interpreting empirical patterns of genetic variation in species with complex genomic structures and demographic histories (Conover and Wendel, 2022).

Mutation-selection balance (MSB) is one mechanism by which genetic variation is maintained by introducing and removing variants at equal rates through mutation and purifying selection, respectively (Crow and Kimura, 1970). MSB quantifies the expected amount of segregating genetic variation at a locus and, by extension, mutation load. This, in turn, facilitates comparisons of the expected variation due to MSB with the variation present in genomic data (e.g., Simmons and Crow, 1977). Although variants can also be maintained through other evolutionary processes such as balancing or linked selection (Marion and Noor, 2023), models of MSB have been used to argue for the maintenance of sex and recombination (Charlesworth, 1990; Chasnov, 2000) and were recently applied to specific classes of mutations (Weghorn et al., 2019) and cancer progression (Persi et al., 2021).

Classical models of MSB find equilibria in deterministic models under an infinite population size limit (Moran, 1977) (see Bürger (2000) for a comprehensive review), but a more general concept of dynamic MSB has been developed for finite populations when considering both deleterious and beneficial mutations (Haigh, 1978; Goyal et al., 2012). The deterministic regime which we adopt here was first considered for a diploid population by Haldane (1927) leading to approximations for the segregating frequency of a mutant allele at MSB as 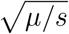 for recessive mutants and *µ/sh* otherwise, where *µ, s*, and *h* are the mutation rate, selection coefficient, and dominance coefficient, respectively (Crow and Kimura, 1970). Ronfort (1999) extended these results to polysomic tetraploids with the approximations 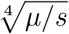 for recessive mutants and *µ/sh*_1_ otherwise where *h*_1_ is the dominance coefficient for an individual with a single mutant copy. Clearly, the mutant allele frequency at mutation-selection balance is sensitive to dominance for both diploids and polyploids. However, the relationship between dominance and selection is not well-understood (Di and Lohmueller, 2024), although many lines of evidence suggest that more deleterious mutations tend to be more recessive (Eyre-Walker and Keightley, 2007; Huber et al., 2018; Kyriazis and Lohmueller, 2024; Zeitler and Gilbert, 2026). Dominance is considerably more complicated in polyploids and, arguably, more important because polyploids are hypothesized to sustain more segregating deleterious variation through the masking of (partially) recessive mutations (Otto and Whitton, 2000). Furthermore, dominance has been shown to shape other evolutionary outcomes in polyploids, including expansion load (Booker and Schrider, 2025) and pseudo-overdominance (Booker and Schrider, 2026). Thus, we focus on exploring how MSB varies with ploidy, mode of inheritance, and dominance.

Here, we address the theoretical gap of mutation-selection balance for polyploids by developing a deterministic mutation-selection model for allotetraploids adapted from the two-locus, two-allele models for diploids (Kimura, 1956; Lewontin and Kojima, 1960; Karlin and McGregor, 1971). We also present a robust approach based on nonlinear dynamics for identifying mutation-selection equilibria under arbitrary dominance models for tetraploids, regardless of whether inheritance is disomic or tetrasomic, and compare this analysis to that for previously developed models of tetraploids with polysomic inheritance patterns and diploids. Using this new framework, we find that when selection is parameterized to act solely on the total dosage of an allele (i.e., selection acts equally on both subgenomes of an allotetraploid), ploidy is the primary determinant of equilibrium allele frequency and mutation load at MSB for both auto- and allotetraploids under the assumption of equal mutation rates and an infinite population size. Furthermore, for completely dominant deleterious mutations with unequal forward and back mutation rates, multiple points of MSB can coexist in both the diploid and tetraploid models. Bistable equilibria have not been previously observed in autotetraploids and could have important implications for evolutionary trajectories. These bistable equilibria arise from a pair of saddle-node bifurcations and are present for a wider range of parameter space in the tetraploid models than in the diploid model. Additionally, for overdominant mutations, only the allotetraploid model exhibits bistability characterized by the fixation of an allele in one subgenome and its loss in the other leading to more heterozygotes and higher fitness. We also compare our numerical estimates for load to existing analytical approximations (Haldane, 1937; Ronfort, 1999). Finally, we investigate the temporal dynamics of how allele frequency and fitness change over time using discrete time simulations. For alleles under purifying selection, we assess the impacts of bistability and bifurcations on the rate of approach to mutation-selection equilibria. Our framework provides quantitative foundations for understanding how ploidy shapes the fundamental balance between mutation and selection, with potential applications to both crop improvement and the prediction of evolutionary trajectories in polyploid populations.

## 2 Model

### 2.1 General Model Description and Notation

Here, we introduce a general mathematical model for the effects of selection and mutation on gamete frequencies in populations with discrete, non-overlapping generations. Specifically, we consider diploids, autotetraploids (tetraploids with polysomic inheritance), and allotetraploids (tetraploids with disomic inheritance), although the modeling approach presented in this section could be applied more generally to any ploidy level with any inheritance pattern. For each generation *t*, we begin with gamete frequencies *g*^*t*^, progress through the processes of random mating, selection, meiosis, and mutation, and conclude with gamete frequencies *g*^*t*+1^ in generation *t* + 1.

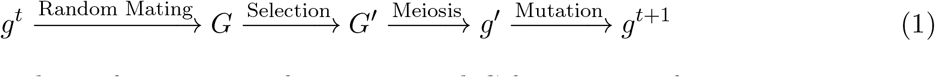

In general, *g* will be used to refer to gamete frequencies and *G* for genotype frequencies.

Under this modeling approach, the gamete frequencies in the next (discrete) generation can be written as an iterative map or difference equation of the form

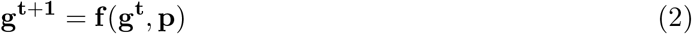

where **g** is a vector of gamete frequencies and **p** is a vector of parameter values for mutation, selection, and dominance.

Then, to find the change in gamete frequency in a single generation, we have

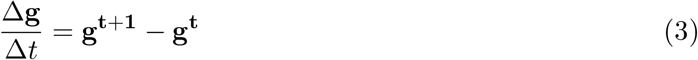

where Δ*t* is one unit of discrete generational time. Because the dynamics are more easily modeled by differential equations, we follow the analysis of Hofbauer (1985) to replace the discrete time difference **g**^**t**+**1**^ − **g**^**t**^ with the continuous time derivative **ġ** = *d***g***/dt*. Then, we have a system of ordinary differential equations (ODEs)

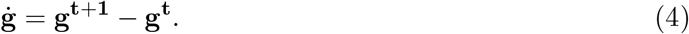

This construction of ODEs is equivalent to that presented for diploids by Bürger (1998).

In the following section, we present the allotetraploid model in detail and define the notation and parameters used throughout the paper. The complete diploid and autotetraploid models are included in the supplementary material, and the differences between these and the allotetraploid model are briefly noted in Section 2.3.

### 2.2 Allotetraploid One-Locus Model

We model a single biallelic locus in an allotetraploid population in which each individual has two distinct subgenomes with independent segregation between them. That is, we do not consider homoeologous exchanges or homoeologous gene conversion (although our general framework allows for such extensions). Our model is conceptually similar to the two-locus diploid model first presented by Kimura (1956) and later extended to include mutation by Karlin and McGregor (1971). Much of the notation is borrowed from Lewontin and Kojima (1960) and Gerard (2023*a*).

#### 2.2.1 Notation and Random Mating

Let **g** be a 2x2 matrix of diploid gamete frequencies with elements *g*_*ij*_ where *i, j* ∈ {0, 1} denotes the number of derived (‘a’) alleles in the subgenomes of each gamete.

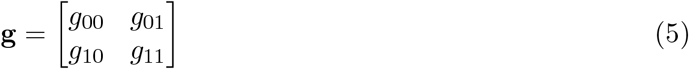

The allele frequencies in each subgenome (i.e., *p*_*a*_, *q*_*a*_, *p*_*b*_, *q*_*b*_) can be determined by computing the marginal frequencies of **g**. Here, *p* denotes the frequency of the ancestral allele (‘A’) and *q* the frequency of the derived allele (‘a’). The allele frequencies in the *a* subgenome are the marginals over the rows of **g**, and the allele frequencies in the *b* subgenome are the marginals over the columns.

Let **G** be a 3x3 matrix of genotype frequencies. Each element, *G*_*ij*_ where *i, j* ∈ {0, 1, 2}, denotes the genotype frequency with *i* and *j* derived alleles in subgenomes *a* and *b*, respectively. Similarly to how the subgenome allele frequencies can be obtained from the marginal distributions of **g**, the genotype frequencies within subgenomes *a* and *b* can be obtained by summing over the rows and columns of **G**, respectively. Under random mating, **G** is defined by a two-dimensional discrete linear convolution of **g**, denoted 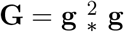 (Gerard, 2023*a*).

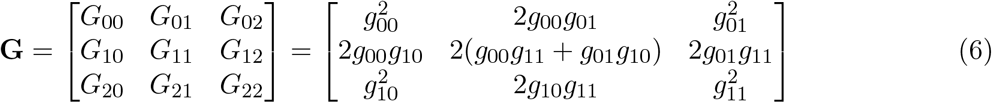

This linear convolution is equivalent to calculating genotype frequencies as the expectation of two independent draws from a multinomial distribution of the gamete frequencies in **g**. The genotype frequencies *G*_*ij*_ are expanded in terms of the gamete frequencies *g*_*ij*_ in the second matrix above and in Column 3 of Table 1.

**Table 1:** Allotetraploid Model Summary for Mutation-Selection Balance.

| Allotetraploid<br>Genotype | Genotype<br>Frequency | Expressed In<br>Gamete<br>Frequencies As | Relative<br>Fitness | Produces Gametes<br>In Proportions<br>$g_{00} : g_{10} : g_{01} : g_{11}$ |
| --- | --- | --- | --- | --- |
| AA \|\| AA | $G_{00}$ | $g_{00}g_{00}$ | 1 | 1 : 0 : 0 : 0 |
| AA \|\| Aa | $G_{01}$ | $2g_{00}g_{01}$ | $1 - sh_1$ | $\frac{1}{2} : 0 : \frac{1}{2} : 0$ |
| Aa \|\| AA | $G_{10}$ | $2g_{00}g_{10}$ | $1 - sh_1$ | $\frac{1}{2} : \frac{1}{2} : 0 : 0$ |
| AA \|\| aa | $G_{02}$ | $g_{01}g_{01}$ | $1 - sh_2$ | 0 : 0 : 1 : 0 |
| aa \|\| AA | $G_{20}$ | $g_{10}g_{10}$ | $1 - sh_2$ | 0 : 1 : 0 : 0 |
| Aa \|\| Aa | $G_{11}$ | $2g_{01}g_{10} + 2g_{00}g_{11}$ | $1 - sh_2$ | $\frac{1}{4} : \frac{1}{4} : \frac{1}{4} : \frac{1}{4}$ |
| Aa \|\| aa | $G_{12}$ | $2g_{01}g_{11}$ | $1 - sh_3$ | 0 : 0 : $\frac{1}{2} : \frac{1}{2}$ |
| aa \|\| Aa | $G_{21}$ | $2g_{10}g_{11}$ | $1 - sh_3$ | 0 : $\frac{1}{2} : 0 : \frac{1}{2}$ |
| aa \|\| aa | $G_{22}$ | $g_{11}g_{11}$ | $1 - s$ | 0 : 0 : 0 : 1 |

### 2.2.2 Allotetraploid Selection

We model selection acting on each tetraploid genotype as a 3x3 matrix of relative fitnesses, **S**, such that each element is the relative fitness of its respective element in **G** (Column 4 of Table 1). We model the strength of selection in allotetraploids as being determined only by the dosage of the selected allele and being independent of the dosage distribution across subgenomes (Booker and Schrider, 2025). Thus, we assume that there is no functional difference between alleles residing on different subgenomes and that variants in each subgenome are seen equally by selection (but note that this model can be easily developed to include subgenome-specific selective effects, or to include homoeologous epistatic dominance, as has been suggested previously (Conover and Wendel, 2022)). In this case, the matrix of relative fitnesses is of the form

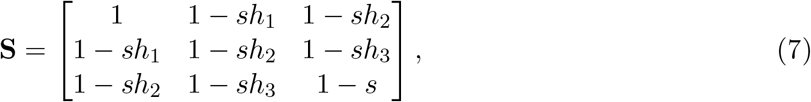

where *s* is the selection coefficient of an allele (here and throughout, we will only explore deleterious mutations) and *h*_1_, *h*_2_, *h*_3_ are the dominance coefficients for individuals with one, two, and three copies of the selected allele, respectively. Note that the sum of the subscripts in **G** corresponds to the subscript of the dominance coefficient in **S** for all heterozygous genotypes. For example, *G*_11_ has two deleterious alleles, one in each subgenome, which corresponds to a dominance coefficient of *h*_2_ in our model of selection.

We define 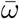 to be the average fitness of the population, calculated as the sum of the elements of **S** with corresponding weights **G**, which is

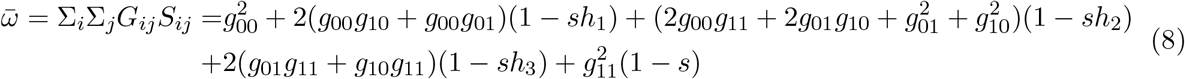

and can be simplified to

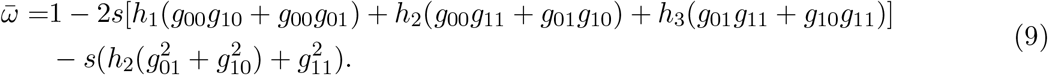

Then, we define **W** to be a matrix of normalized fitnesses, with 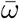 as the standard normalizing factor. So, 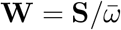 gives

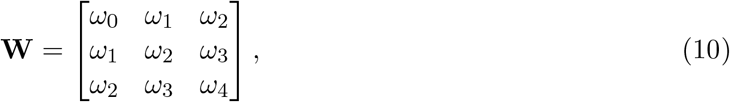

where *ω*_*i*_ is the normalized fitness effect for a genotype with *i* derived alleles. Finally, the normalized genotype frequencies after selection, **G**^*′*^, are

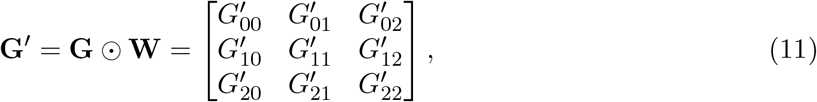

where ⊙ denotes the Hadamard product which is an element-wise multiplication of two matrices.

For example, 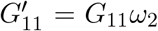. The *ij*^*th*^ element of **G**^*′*^ is simply

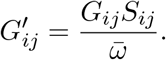

### 2.2.3 Allotetraploid Meiosis

To model gamete segregation for a disomic allotetraploid, we use the following system of equations

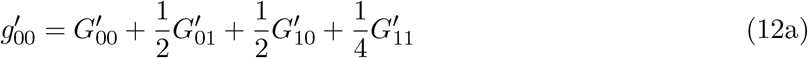

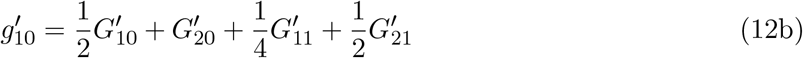

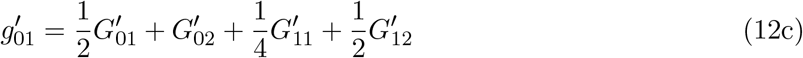

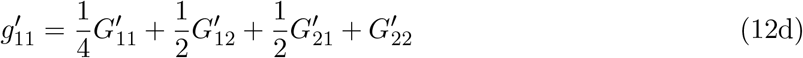

where the coefficients for each term are the proportions of that gamete generated from the given allotetraploid genotype. For example, the coefficient for 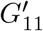 is 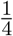 for all gametes because the *G*_11_ genotype produces one of each gamete type, so the proportion of each gamete is one in four. These segregation frequencies are listed in Column 5 of Table 1 and are equivalent to those for a diploid two-allele, two-locus model (Li, 1972). The four simplified equations above are equivalent to equations 10.a - 10.d studied by Lewontin and Kojima (1960) when the recombination rate, *R*, is 0.5 (i.e., loci are on separate chromosomes, as is the case for homoeologous loci in an allopolyploid).

#### 2.2.4 Allotetraploid Mutation

We define the rate of forward mutation from the ancestral to derived state as *µ* and the rate of back mutation as *ν*. We assume that the mutation rates are equal between subgenomes and that mutation acts only on the gametes after meiotic segregation. The following set of equations model the transition probabilities between types of gametes in terms of *µ* and *v*.

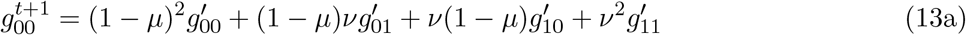

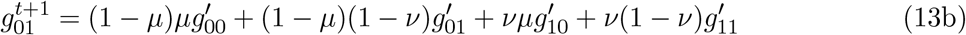

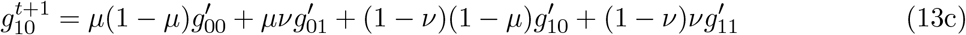

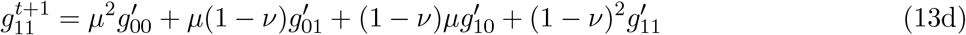

For example, in (13a), 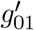 will result in a 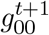 gamete if two specific events occur i) no forward mutation occurs in the *a* subgenome (hence the 1 − *µ* factor) and ii) back mutation does occur in the *b* subgenome (hence the *v* factor). The other mutation terms are similarly ordered.

#### 2.2.5 Allotetraploid Changes in Gamete Frequencies

Combining the steps of random mating, selection, meiosis, and mutation described above, we model the change in gamete frequencies between subsequent generations by subtracting a 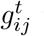 term from (13) to obtain the following system of ODEs for allotetraploids:

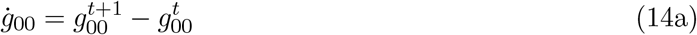

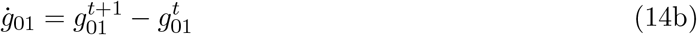

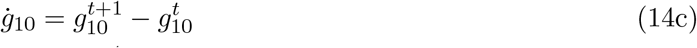

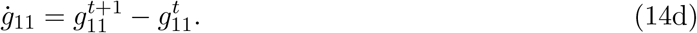

This system of equations can then be reduced to a system of three equations in three variables by progressive substitution of equations (6), (11), (12), and (13) into equation (14) and finally substituting *g*_10_ = 1 − *g*_00_ − *g*_01_ − *g*_11_ into the resulting system and removing the *ġ*_10_ equation to account for the linear constraint that gamete frequencies sum to one. The choice of which gamete frequency and corresponding equation to remove is arbitrary, but the removal of *g*_10_ and *ġ*_10_ is reflected in our corresponding numerical analysis.

### 2.3 Comparison to Diploid and Autotetraploid Models

The notation and model structure for both diploids and autotetraploids is conceptually similar to that presented above for allotetraploids. The primary difference is that diploids and autotetraploids lack any subgenomic structure and, thus, gamete and genotype frequencies **g** and **G** are modeled as vectors instead of matrices. By extension, selection is also modeled as a vector of fitnesses instead of as a matrix. Importantly, the diploid and autotetraploid models use slightly different notation for gametes and genotypes; instead of an *ij* subscript which delineates between subgenomes, both models use a single *i* subscript for the total number of derived alleles in a gamete or genotype. Additionally, while traditional diploid models focus on changes in allele frequencies, we instead refer to gamete frequencies for semantic and notational consistency across all three models. Gamete and allele frequencies are equivalent for the diploid model, and, when allele frequencies are of interest, we use the gamete frequencies in the tetraploid models to calculate allele frequency as a summary statistic. To make the distinction between alleles and gametes clear, we use the notation from Lewontin and Kojima (1960) and other two-locus, two-allele diploid models. For comparison with Table 1, general summaries of the diploid and autotetraploid models are in Tables S1 and S2.

## 3 Results

### 3.1 Efficacy of Selection and Bistable Equilibria

Here, we consider how mutation, selection, and dominance collectively shape equilibrium points of mutation-selection balance (MSB); compare these equilibria between the diploid and tetraploid models; and investigate how the location of these equilibria shift as parameters vary.

We begin by considering how points of MSB shift as the strength of selection *s* varies when forward and backward mutation rates are equal (*µ* = *v*) (Fig. 1). In this case, the auto- and allotetraploids have nearly identical equilibrium allele frequencies and mutation loads (Fig. 1). Notably, the two tetraploid models show very slight differences at MSB across dominance classes with the differences being O(*µ*) and O(*µ*^2^) for allele frequency and mutation load, respectively (Figs. S2, S3). As expected, the frequency of the deleterious allele decreases as *s* increases, but the response to selection is not equal between the tetraploid and diploid models (Fig. 1A-E). The tetraploid populations generally experience a smaller change in allele frequency for the same change in the strength of selection. Additionally, for weak selection (approximately *s < µ*), the equilibria arise from a balance between mutational pressures with the approximate equilibrium given by *q* ≈ *µ/*(*µ* + *v*) (Crow and Kimura, 1970). In this case where *µ* = *v*, it is straightforward to see that the expected allele frequency for a neutral or nearly neutral allele (*s* ≪ 1) is *q* = 0.5 (Fig. 1A-E).

**Figure 1:**
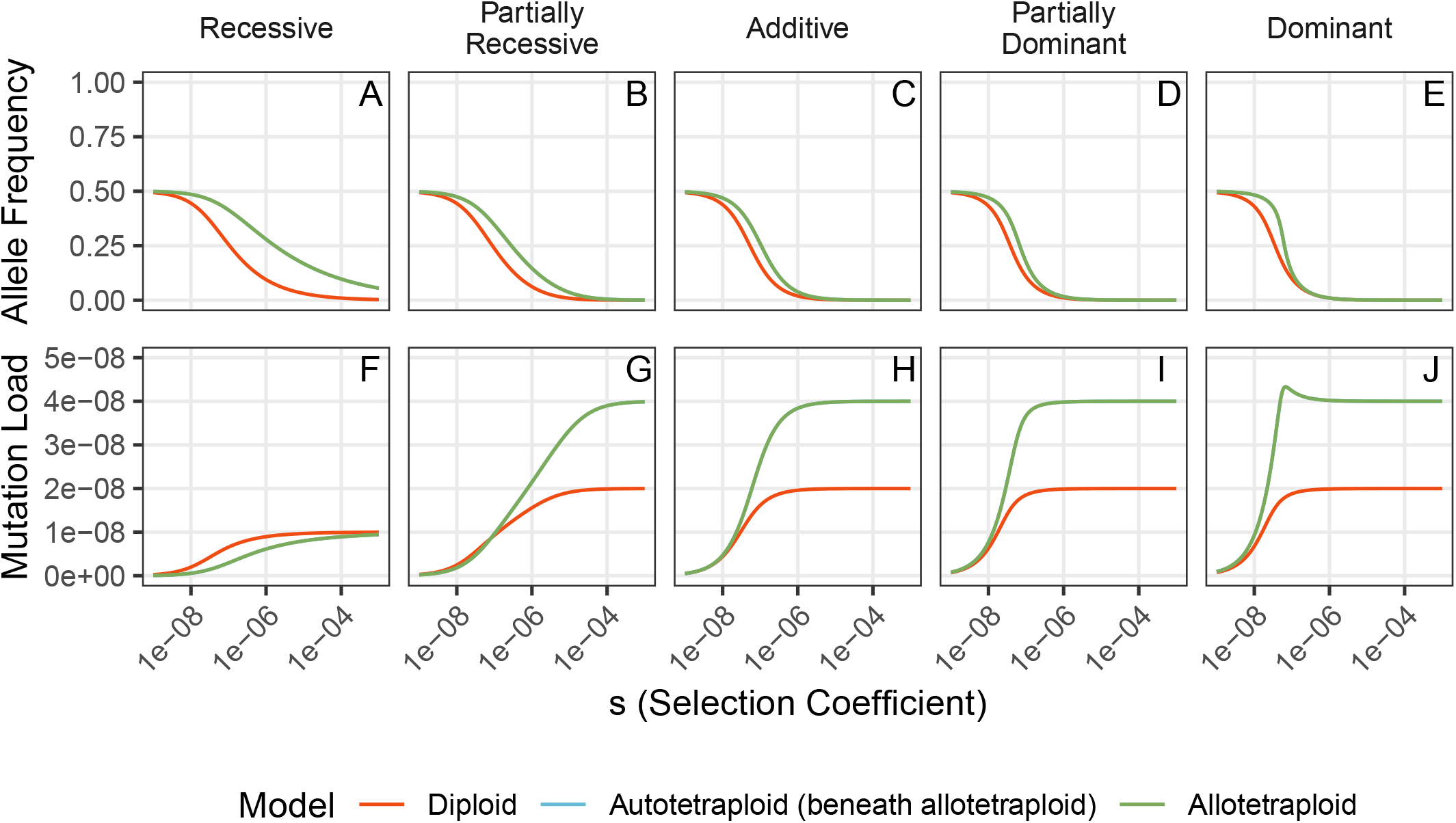
Mutation-selection equilibria for equal mutation rates. Allele frequency and mutation load at equilibrium for 10^*−*9^ *< s <* 10^*−*3^ are shown across dominance regimes. For all panels, *µ* = *v* = 10^*−*8^. Dominance coefficients for the partially recessive column are *h*_1_ = 0.02, *h* = *h*_2_ = 0.1, and *h*_3_ = 0.4 and for the partially dominant column are *h*_1_ = 0.6, *h* = *h*_2_ = 0.9, and *h*_3_ = 0.98.

Furthermore, both the auto- and allotetraploids have a higher proportion of deleterious alleles segregating at MSB compared to diploids facing the same mutation and selection pressures, regardless of dominance; selection is less efficacious at purging deleterious alleles at higher ploidies. However, these higher allele frequencies do not necessarily imply a lower fitness. This is because tetraploids can mask the effects of recessive and partially recessive deleterious mutations; specifically, in line with theoretical predictions (Otto and Whitton, 2000), we find that tetraploids have a mutation load lower than the expected *L* = 4*µ* for partially recessive mutations provided that *h*_1_ *< h/*2 (Fig. 1G). In the fully recessive case, we find that the tetraploid populations are more fit than the diploid populations for all *s* when measured by mutation load *L* (Fig. 1F). Remarkably, under the special case of equal mutation rates, *L* is up to twice as large for diploids as for tetraploids at intermediate strengths of selection for fully recessive mutations (Fig. 1F). Haldane (1937) first analytically calculated *L* = *µ* in the recessive case for diploids, and Ronfort (1999) extended this result to higher ploidy, which agrees with our numerical estimates of *L* for large *s*. Note that because our model includes back mutation, our numerical estimates of *L* disagree with the analytical approximation of Ronfort (1999) for small *s*. For the cases with any dominance, load is approximately twice as large for the tetraploid populations which reflects that there are twice as many loci/chromosomes to mutate (Fig. 1G-J). For non-recessive dominance, Otto and Whitton (2000) calculated *L* ≈ *kµ* for a population of *k*-ploid individuals, which generally aligns with our results (but note the deviation for partially recessive mutants). Surprisingly, we find a non-monotonic relationship between *s* and *L* for dominant mutations in the tetraploids near *s* = 10^*−*7^ (Fig. 1J). This unexpected increase in load can be attributed to non-zero panmictic disequilibrium and the genotype distribution at MSB, as there are more individuals in the population with a single copy of the deleterious allele than expected under HWE when *s* ≈ 10^*−*7^, compared to when *s <* 10^*−*7^ or *s >* 10^*−*7^ (Sec. S2.1, Fig. S4).

We now turn our attention to a case of biased mutation rates such that *µ > v*, which, although not directly considered here, includes the classical case without back mutation (i.e., *v* = 0) (Fig. 2). In this region of parameter space, most qualitative results observed under the equal mutation rate regime still apply. For example, the deleterious allele has a higher segregating frequency at MSB in tetraploids, and this frequency decreases as the strength of selection increases. Additionally, the same approximations for mutation load hold for stronger strengths of selection, and for *s < µ* the populations exist in a mutational equilibrium at *q* = *µ/*(*µ* + *v*), which for *µ* = 2 × 10^*−*8^ and *v* = 10^*−*9^ gives *q* ≈ 0.95. For the recessive and additive mutations, there is still a single, globally stable equilibrium which all populations will evolve toward. However, for dominant mutations, we find a more interesting result; for some values of *s*, there are two stable fixed points (Fig. 2C). This is termed bistability and was originally reported in diploids by Haldane (1927) and can be understood by decomposing the differential equation into separate components due to mutation and selection (Fig. S5). In this case, the equilibrium at a high allele frequency is due to balance between the forward and backward mutation pressures (with relatively weak contributions from selection) while the equilibrium at a low allele frequency is due to a balance between variation being introduced by the forward mutation process and removed by purifying selection (with relatively weak contributions from back mutation). For these dominant mutations, the presence of bistability indicates that populations facing the same selective and mutation pressures, but with different initial allele frequencies, may evolve toward different equilibria (Fig. S6). Thus, a population can evolve to have lower fitness and a higher deleterious allele frequency, despite facing the pressure of purifying selection. For example, a population just above the unstable equilibrium (dashed line) will evolve up toward a stable fixed point near *q* ≈ 0.95 (Fig. 2C). The mutation load *L* for these high allele frequency equilibria are not independent of the strength of selection as predicted by classical MSB theory (Haldane, 1937) and instead *L* increases linearly with *s*. For the low allele frequency equilibria maintained by a balance between selective and forward mutational pressures, *L* is independent of *s* and approximately equal to 2*µ* and 4*µ* for the diploids and tetraploids, respectively (Fig. 2F).

**Figure 2:**
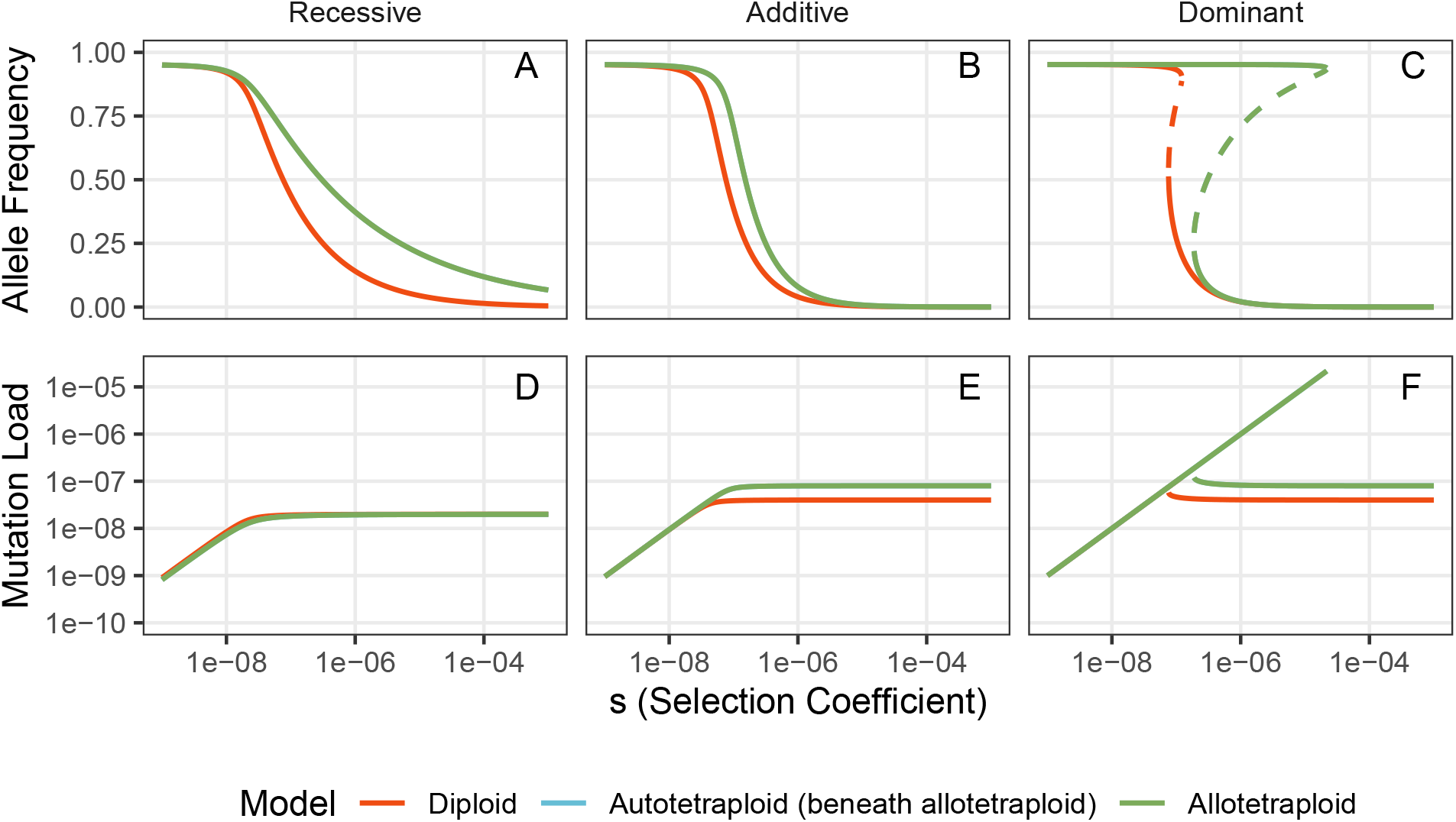
Mutation-selection equilibria for biased mutation rates. Allele frequency and mutation load at equilibrium for 10^*−*9^ *< s <* 10^*−*3^ are shown across dominance regimes. For all panels, *µ* = 2 × 10^*−*8^ and *v* = 10^*−*9^. The solid lines represent stable equilibria. The dashed lines in (C) represent unstable equilibria. In (F), load is only shown for the stable equilibria.

Finally, we explore how equilibria shift in response to changes in the strength of selection for an overdominant deleterious mutation. Specifically, we model overdominance by setting *h* = *h*_1_ = *h*_2_ = *h*_3_ = −0.05, which, for diploids, is well within estimated values of *h* for quantitative traits exhibiting overdominance (e.g., estimates of *h* are −0.175 for the classical example of sickle cell trait in human populations with exposure to malaria (Allison, 1964; Hartl, 2020)). Our model of overdominant selection is similar to the model of stabilizing selection acting on a single locus originally studied by Robertson (1956). Across each of our diploid and tetraploid models, when *s < µ*, the equilibrium allele frequency is determined by a balance between the forward and backward mutation rates (Fig. 3A). As *s* increases, the equilibrium allele frequency decreases and segregation load increases linearly for the diploid and autotetraploid populations (Fig. 3A,B). This is a result of our model of selection in which individuals homozygous for the derived allele still have a small fitness cost for the population (i.e., derived and ancestral homozygotes do not have equal fitness), which pushes equilibria below *q* = 0.5. The equilibrium allele frequency arises from a balance between the fitness benefit of heterozygotes and the fitness consequence of derived homozygotes. The allotetraploid model exhibits qualitatively different dynamics characterized by bistability in the overdominant regime with an unstable branch and two stable branches emerging from a pitchfork bifurcation at *s* ≈ 10^*−*6^ (Fig. 3C). The unstable branch is restricted to the line where *q*_*a*_ = *q*_*b*_ and mirrors the equilibrium of the autotetraploid model when measuring either average allele frequency between subgenomes or load (Fig. 3). As the strength of selection increases, the stable branches in the allotetraploid evolve toward fixation in one subgenome and loss in the other which results in substantial gametic heterozygosity and, in turn, high levels of genotypic heterozygosity (Fig. 3C). Compared to the diploid and autotetraploid models in which load increases with *s*, the stable equilibria of the allotetraploid maintain constant segregation load proportional to the influx of new mutations each generation (Fig. 3B). These results suggest that the ‘fixed heterozygosity’ that is typical of allotetraploid genomes may not only improve fitness by masking the fitness effect of recessive deleterious alleles, but can also effectively decouple the selective consequences of overdominant alleles into independently segregating subgenomes.

**Figure 3:**
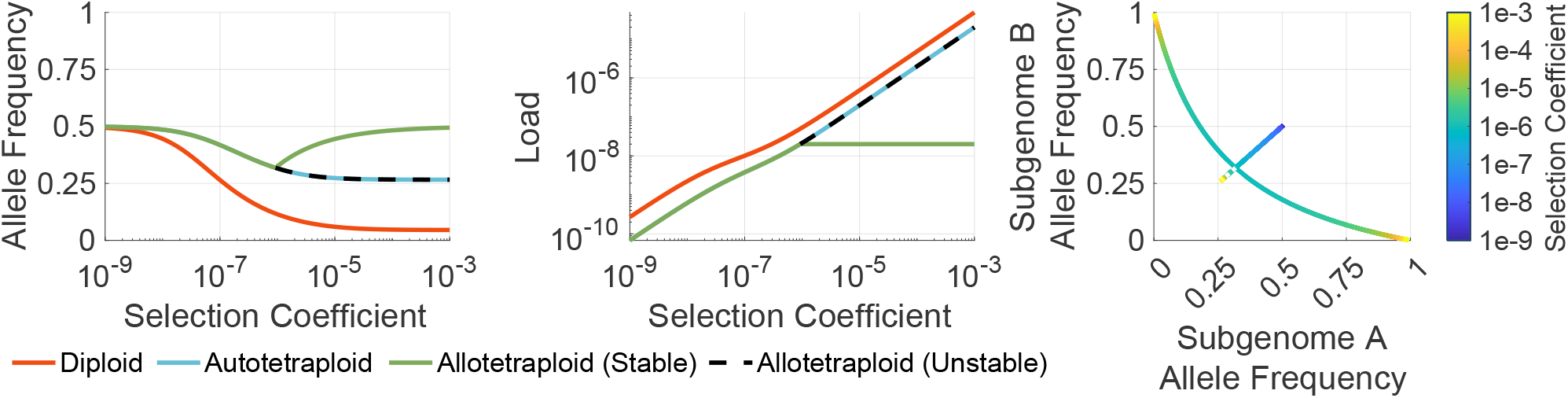
Mutation-selection equilibria for overdominant mutations. Allele frequency and segregation load are shown for 10^*−*9^ *< s <* 10^*−*3^, *µ* = *v* = 10^*−*8^, and *h* = *h*_1_ = *h*_2_ = *h*_3_ = −0.05. In (A), the allele frequency for allotetraploids is the average across subgenomes. In (C), the pitchfork bifurcation for allotetraploids is shown in subgenome-specific allele frequency space with dashed lines indicating unstable equilibria.

### 3.2 Efficiency of Selection and Temporal Dynamics

To investigate the deterministic temporal dynamics as populations approach MSB equilibrium, we developed a discrete time simulation that tracks the evolution of gamete frequencies for a specified number of generations and arbitrary initial conditions, but still assuming an infinite population size. Specifically, we model populations in which the proportion of gametes that are involved in creating generation *t* + 1 is exactly equal to the proportion at which they are formed in generation *t*, rather than randomly sampling gametes each generation as would be done in a stochastic modeling approach. We compare the results of these simulations between the diploid and tetraploid populations, with a focus on the impact that bifurcations and bistability described above have on distorting the temporal dynamics. We first consider a case with balanced mutation rates that are weak relative to the strength of selection (Fig. 4).

**Figure 4:**
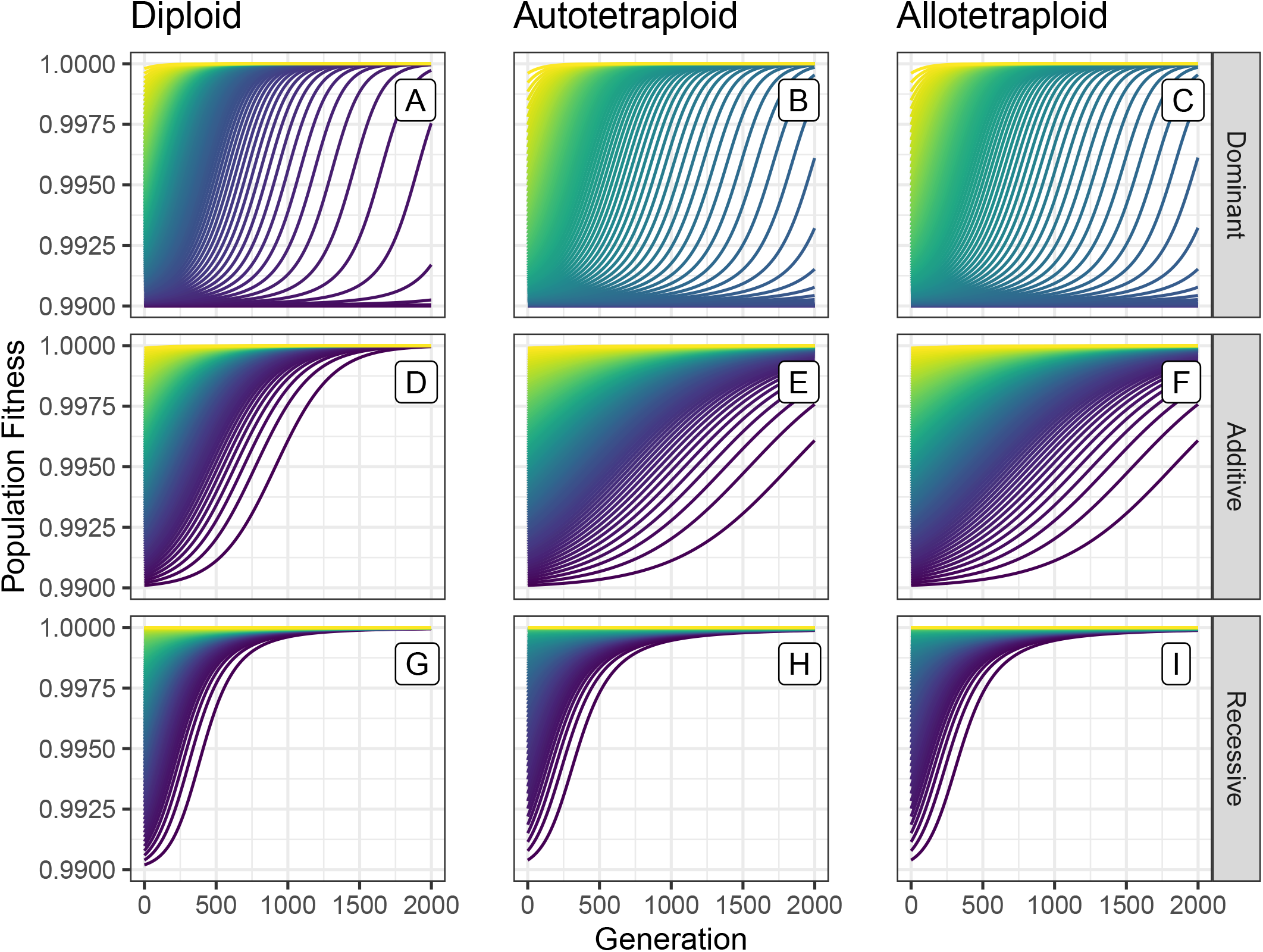
Fitness trajectories across models and dominance for weak, unbiased mutation. Trajectories of population fitness over time for *s* = .01, *µ* = 10^*−*8^, and *v* = 10^*−*8^. Each trajectory shows the evolution of a single simulation beginning in Hardy-Weinberg Equilibrium with an initial allele frequency between 0.01 and 0.99 incremented by 0.01. The coloration denotes the initial allele frequency when the simulation started (i.e., at *t* = 0). For allotetraploids, the same initial allele frequency was used in both subgenomes.

Here, we consider trajectories of both population fitness (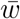, Fig. 4) and allele frequency (*q*, Fig. S7) as summaries of the simulations to explore how dominance alters temporal dynamics over time. For population fitness, we find that diploids, autotetraploids, and allotetraploids show the same general pattern: population fitness approaches the equilibrium value more quickly for recessive variants than for additive or dominant ones (Fig. 4). However, this does not directly correspond to a more efficient purging of the deleterious allele from the population (Fig. S7). Instead, additive deleterious alleles are most efficiently removed by natural selection, but each remaining copy of the allele still has an impact on the fitness of the population (whereas a recessive allele only affects the population fitness if it is present in a homozygous state). The masking of (partially) recessive deleterious mutations in tetraploids also results in identical rates of fitness increase between the diploid and tetraploid populations (Fig. 4G-I) (akin to the effect of masking on mutation load seen in Fig. 1F and Fig. 2D), even though the allele frequency at MSB is much higher in the tetraploids than in diploids (Fig. S7). Thus, for recessive mutations, purifying selection is equally effective and efficient in the diploid and tetraploid models, but for additive and dominant mutations, the tetraploids evolve more slowly as measured by both fitness and allele frequency, which reflects theoretical predictions of the response to selection in polyploids (Ronfort, 1999; Otto and Whitton, 2000).

Next, we consider a case with biased mutation rates (specifically 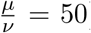), in which each rate is comparable to the strength of selection (Fig. 5). We note that, to improve the speed of these simulations, we use artificially high mutation rates and strongly deleterious selection coefficients; however, rescaling these mutation rates and selection coefficient by a constant produces qualitatively similar trajectories (Fig. S9). Due to masking effects in polyploids, the diploids and tetraploids evolve at an equivalent rate for recessive mutations when measuring population fitness (Fig. 5G-I). In the biased mutation rate regime, allele frequency decreases most quickly for recessive mutations because bifurcations near the additive and dominant parameter space slow the temporal dynamics (Fig. S10). As a result, the diploid and tetraploid populations evolve on similar time scales for dominant mutations (as opposed to the dominant mutations evolving relatively faster in the diploid population under equal mutation rates).

**Figure 5:**
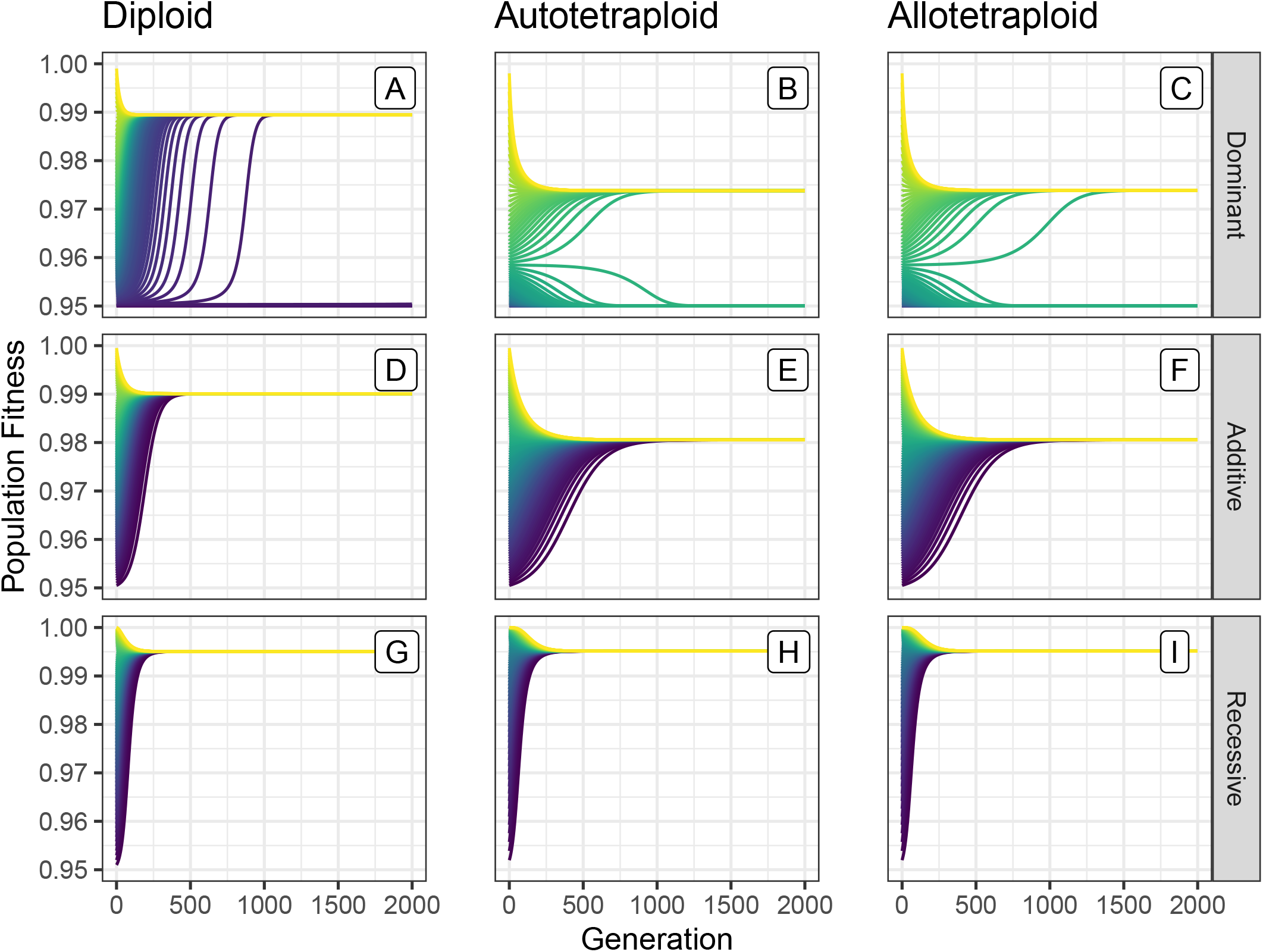
Fitness trajectories across models and dominance for strong, biased mutation. Trajectories of population fitness over time for *s* = .05, *µ* = 5×10^*−*3^, and *v* = 10^*−*4^. Each trajectory shows the evolution of a single simulation beginning in Hardy-Weinberg Equilibrium with an initial allele frequency between 0.01 to 0.99 incremented by 0.01. The coloration denotes the initial allele frequency when the simulation started (i.e., at *t* = 0).

Finally, we consider the evolutionary trajectories of an overdominant deleterious mutation across models. Across models, segregation load decreases as the populations evolve toward equilibrium, although this decrease in load occurs to varying extents and on differing time scales (Fig. 6). Note that for overdominant mutations we plot the segregation load defined in Eqn. 15 as opposed to fitness since in the overdominant case there is not a one-to-one correspondence between the two (Fig. S12 shows the corresponding fitness trajectories). The diploid populations rapidly approach equilibrium values of allele frequency and segregation load regardless of the starting allele frequency (Fig. S13A). Similarly, autotetraploid populations with an initial allele frequency greater than the equilibrium frequency quickly approach equilibrium. However, this is not the case for autotetraploid populations starting with an initial allele frequency lower than the equilibrium frequency (i.e., the yellow trajectories shown in Figs. 6B and S13B). In this case, allele frequency stays considerably below the equilibrium value for the duration of the simulations contributing to elevated segregation load (Fig. 6B). The temporal dynamics are more dramatically slowed for the allotetraploid populations which take more than 100,000 generations to reach equilibrium (Fig. 6C). With the exception of the trajectory evolving along the unstable branch *q*_*a*_ = *q*_*b*_, the allotetraploid populations evolve to have less segregation load after a long transient period through complementary fixation and loss of the derived allele in separate subgenomes (Fig. S14).

**Figure 6:**
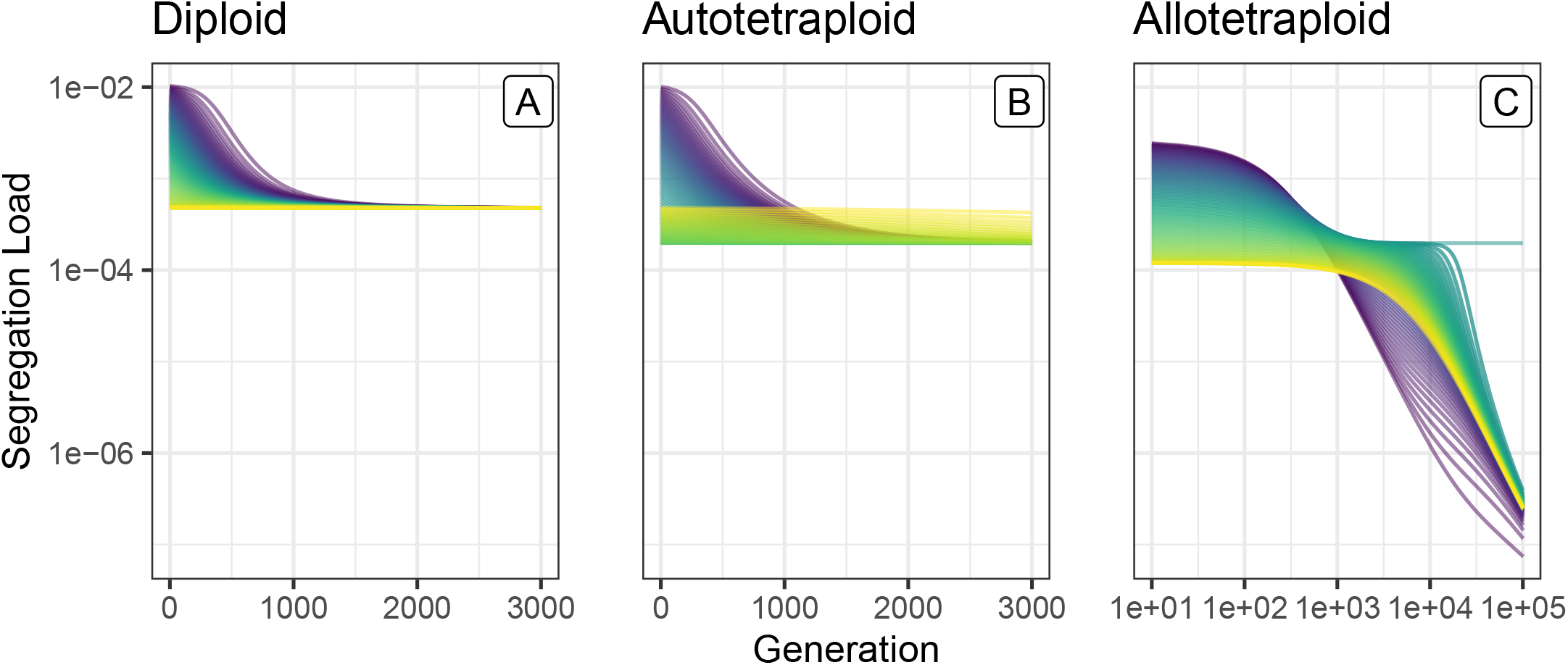
Load trajectories across models for overdominant selection. Trajectories of segregation load over time for *s* = 0.01, *h* = *h*_1_ = *h*_2_ = *h*_3_ = −0.05, and *µ* = *v* = 10^*−*8^. Each trajectory shows the evolution of a single simulation beginning in Hardy-Weinberg Equilibrium with coloration denoting the initial allele frequency when the simulation started (i.e., at *t* = 0). For the diploids and autotetraploids, initial allele frequencies were chosen in increments of 0.01 ranging from 0.01 to 0.99. For the allotetraploid simulations, initial allele frequencies in one subgenome were incremented by 0.01 ranging from 0.01 to 0.99 and were held fixed at 0.5 in the second subgenome. Note that the allotetraploid panel has a different horizontal scale.

## 4 Discussion

We have developed a nonlinear dynamics approach to construct the first comprehensive mathematical framework for comparing mutation-selection dynamics across diploid, autotetraploid, and allotetraploid systems with arbitrary dominance coefficients, addressing a critical theoretical gap. Our deterministic models provide quantitative foundations for understanding how ploidy level fundamentally reshapes evolutionary equilibria by extending classical one- and two-locus biallelic diploid models into tetraploids with disomic and tetrasomic inheritance patterns. Our framework predicts that for most modes of selection auto- and allotetraploids exhibit remarkably similar evolutionary trajectories despite their distinct meiotic behaviors, with both types showing higher segregating allele frequencies compared to diploids at mutation-selection balance (Fig. 1). Additionally, for completely recessive mutations, both tetraploid models are at least as fit as and evolve at equivalent rates as diploid populations (Figs. 1, 4). This suggests that, under the model of mutation and selection we consider, ploidy — and the associated masking of deleterious alleles — may be more important than the specific mode of inheritance. Perhaps most interestingly, we found bistability across all three models for dominant deleterious mutations with biased mutation rates (Fig. 2). Beyond altering the equilibrium structure, this bistability also has substantial impacts on the temporal dynamics of evolution under biased mutation rates with recessive mutations evolving faster than additive or dominant ones (Fig. 5). Finally, for overdominant mutations, we find a distinct equilibrium structure and bistability in the allotetraploid model, which leads to lower segregation load relative to the diploid and autotetraploid models (Fig. 3). The bistability and bifurcation in the overdominant allotetraploid model also slows the temporal dynamics and affects the approach to equilibrium (Fig. 6).

However, the near equivalence of equilibria for autotetraploids and allotetraploids is predicated on our idealized model which considers a single locus in a population with an infinite size and disregards any selective differences between subgenomes in allotetraploids. In empirical populations, established allopolyploids often exhibit subgenome dominance, where homoeologous genes from one subgenome are expressed at universally higher (or lower) levels on average (Edger et al., 2017; Bird et al., 2018), which will influence the degree to which selection will act on variants within each subgenome equally. Furthermore, our model does not consider possible complications due to multivalent formation in polyploids such as homoeologous exchanges (Mason and Wendel, 2020; Wang et al., 2024) or double reduction (Haldane, 1930; Butruille and Boiteux, 2000; Gerard, 2023*b*), although our flexible framework could be extended to explore these dynamics in future work.

### 4.1 Mutation-Selection Balance and Bistable Equilibria

To illustrate how bistability may impact a population’s trajectory, we consider a dominant deleterious mutation’s evolution in a tetraploid population undergoing a severe bottleneck upon formation, as is expected for many neopolyploids (Ramsey and Schemske, 2002). In this case, the initial allele frequency in the neotetraploid will be drawn from a distribution centered on the diploid allele frequency at MSB. Then, in the initial generations while the neotetraploid population remains small, genetic drift will be the primary determinant of the allele’s evolutionary trajectory allowing alleles to drift to widely varying frequencies including loss and fixation in the population. As the neotetraploid population emerges from the bottleneck and population size increases, the dynamics of allele frequency change will more closely match our model and some alleles that began at the low, diploid MSB frequency will evolve toward the high allele frequency and load MSB equilibrium in the tetraploids. Concretely, for a dominant mutation with *s* = 10^*−*6^, the initial frequency in the neotetraploid will be approximately 0.02 on average. Some trajectories will drift up to a frequency above the unstable manifold for tetraploids shown in Fig. 2C causing the allele to evolve up to the equilibrium with an allele frequency of approximately 0.95.

The band of *s* values for which bistability exists is notably larger for tetraploids than for diploids (Fig. 2). Thus, we expect the initial allele frequency — as determined by the interplay of drift, mutation, selection, and founder effects during the bottleneck — to play a more substantial role in the long-term evolutionary trajectory of tetraploids than for diploids. This may contribute to a higher mutation load following the founding bottleneck, even with the increased masking of recessive or partially recessive mutants and reduced inbreeding depression during a bottleneck in autotetraploids (Layman and Busch, 2018; Vlček et al., 2025). Additionally, recent work has focused on the interactions of the increased masking of strongly deleterious, recessive alleles with low recombination rates in small population sizes, revealing that pseudo-overdominance is more prevalent among autotetraploid populations than diploid ones (Booker and Schrider, 2026).

In addition to its implications for neopolyploids, bistability is also relevant to other contexts such as linked selection. Bistability may impact the ability of purifying selection to effectively purge deleterious alleles linked to beneficial mutations because these deleterious alleles may reach a high segregating frequency due to linked selection. Once the linked (dominant) deleterious allele crosses the threshold of the unstable equilibrium, it will remain at a high frequency near the mutational equilibrium and be unable to be removed by purifying selection. Thus, bistability, in conjunction with other evolutionary processes such as bottlenecks and linked selection, has the potential to alter a population’s long-term evolution.

We emphasize that these dynamics are observed in our infinite population size model with recurrent mutation and, thus, only account for the average change in allele frequency in each generation. Variation about this mean change can be caused by various stochasticities in both auto- and allotetraploids; for example, genetic drift leads to differing fixation dynamics in allotetraploids and autotetraploids (Booker and Schrider, 2025) and environmental fluctuations have been shown to have substantial impacts on successful polyploid establishment (Gaynor et al., 2025). The stochastic effects of genetic drift have been considered in both diffusion (Blischak et al., 2023) and coalescent frameworks (Arnold et al., 2012; Roux et al., 2023), although without selection in both cases. Future work incorporating stochastic genetic drift and selection will be especially important for understanding how finite population sizes may interact with the bistable dynamics we find due to the interplay of selection and mutation. In particular, in a finite population size model, the unstable manifold which separates the two stable equilibria will become increasingly blurred as the population size decreases and the effects of genetic drift dominate the impact of mutation and selection.

Even though dominant deleterious mutations are thought to be rare (Eyre-Walker and Keightley, 2007), the bistable dynamics described above for dominant deleterious mutations create important alternative evolutionary trajectories where populations with identical selective pressures but different initial allele frequencies evolve toward dramatically different equilibria — a phenomenon particularly pronounced in polyploids due to their wider parameter ranges for maintaining bistability compared to diploids. Both higher allele frequencies at MSB and bistability may contribute to increased genetic variation in polyploids (Soltis and Soltis, 2000) which has been hypothesized to play a substantial role in the success and prevalence of polyploids among domesticated crops (Salman-Minkov et al., 2016; Qi et al., 2021; McKibben and Barker, 2026). Because of its potential impact on genetic variation, bistability may impact range expansion which is thought to be facilitated by increased plasticity and genetic variance at higher ploidy levels (Ramsey, 2011), although empirical evidence for this hypothesis is mixed (Martin and Husband, 2009; Baniaga et al., 2020; Meirmans and Kolář, 2025) and simulations suggest that inheritance patterns (i.e., disomic vs. tetrasomic) play a crucial role in range expansion and polyploid establishment in ecological niches (Booker and Schrider, 2025). Beyond bistability, we also demonstrate the importance that dominance has on mediating fitness differences between diploids and tetraploids, especially given the potential to mask recessive or partially recessive deleterious mutations. Overall, coupled with complex connections between dominance and selection as well as challenges in estimating the genome-wide distribution of dominance coefficients (Huber et al., 2018; Kyriazis and Lohmueller, 2024; Zeitler and Gilbert, 2026) (see Di and Lohmueller (2024) for a recent review), the presence of both bistability and masking effects in our model makes understanding the joint distribution of dominance and selection coefficients even more important for predicting the evolution of polyploid populations over time.

Finally, we note that our model of overdominant selection is closely related to classical models of stabilizing selection (Robertson, 1956). Specifically, for the allotetraploid model, we parameterize stabilizing selection on the overall genotype or gene expression across subgenomes (as opposed to overdominant selection on each subgenome independently). Under this model, we observe emergent underdominant or disruptive selection in each subgenome such that the allele is lost in the subgenome with lower initial allele frequency and fixed in the other subgenome (Figs. 3 and S14). These emergent dynamics mirror classical results in which models of stabilizing selection for polygenic traits lead to underdominant dynamics at individual loci (Lande, 1976).

### 4.2 Bistability and Temporal Dynamics

For small, equal mutation rates, the diploid and tetraploid populations evolve as expected; masking effects account for equal rates of evolution for recessive mutations and the diploids evolve more quickly for additive and dominant mutations (Fig. 4). Alternatively, for larger, biased mutation rates which admit bistability, the dynamics are notably different. In particular, for both additive and dominant mutations, the diploids and tetraploids evolve on similar time scales (Fig. 5). We also see similar temporal distortions for overdominant mutations in allotetraploids (Figs. 6 and S13). This is due to the presence of bistability and associated bifurcations that distort the temporal dynamics over the transient period before the population reaches an equilibrium. In particular, because the rates of change are all simultaneously zero at an equilibrium, the change in allele frequency per generation for a population near an equilibrium is also very small. Thus, the populations which start near the unstable equilibrium initially evolve very slowly. In fact, the bistability and bifurcations present in the dominant case are also likely responsible for the relatively slower dynamics for additive mutations (compared to recessive ones).

For example, consider a population with partial dominance (not shown here) very near a bifurcation so that for dominance coefficients closer to additivity there is a single stable equilibrium and for dominance coefficients closer to full dominance there are two stable equilibria. In this case, the rates of change are very nearly zero at the emerging bifurcation (because the bifurcation point is an equilibrium itself, the rates of change are necessarily near zero). So, for additive mutations, a population starting at a high allele frequency must pass through the ‘ghost’ left behind by the bifurcation resulting in a long transient period before reaching the MSB equilibrium. This is one example of critical slowing down near a bifurcation which is a well-documented phenomenon in biological models (Van Nes and Scheffer, 2007; Chisholm and Filotas, 2009; Dakos and Bascompte, 2014) and in dynamical systems more broadly (Mandel and Erneux, 1987; Tredicce et al., 2004; Strogatz, 2018).

### 4.3 Conclusion

Our unified mathematical framework provides the first comprehensive comparison of mutation-selection dynamics across diploid, autotetraploid, and allotetraploid systems for a given model of mutation and selection. Within our deterministic, single-locus model, auto- and allotetraploid systems exhibit remarkably similar evolutionary trajectories and mutation-selection equilibria. Most notably, our analysis reveals widespread bistability in polyploid systems which in turn distorts the associated temporal dynamics. While these findings emerge from simplified theoretical conditions, they provide important foundations for understanding when and how genome duplication alters evolutionary processes.

Our flexible framework can be extended to more complex models, such as unequal mutation rates between subgenomes, or cases where selection acts unequally across subgenomes of allotetraploids; unequal selection may arise as a result of homoeolog expression biases, neofunctionalization, or sub-functionalization, among other mechanisms (Adams et al., 2003; Des Marais and Rausher, 2008). Future extensions of our framework could incorporate linkage effects, environmental variation, additional modes of selection, or homoeologous exchange (Mason and Wendel, 2020) and may reveal further complex dynamics including limit cycles, Hopf bifurcations, or additional fixed points not seen under our model of selection but that have previously been documented in two-locus, biallelic diploid models similar to our model for allotetraploids (Akin, 1982; Pontz et al., 2018). Furthermore, while we adopt distinct models of disomic and polysomic inheritance, other frameworks such as those proposed by Meirmans and Van Tienderen (2013) and Blischak et al. (2023) consider a continuum from autopolyploidy and polysomic inheritance to allopolyploidy and disomic inheritance.

The substantial theoretical gap in polyploid population genetics relative to diploids (Meirmans et al., 2018) has limited our ability to predict and interpret evolutionary patterns in polyploid populations. This theory is critical for identifying the action of mutation, selection, and drift in evolution, especially among many complicating processes in polyploids such as double reduction, dosage effects, and biased fractionation. Our nonlinear dynamics framework begins to address this need and provides a foundation for analyzing deterministic mutation-selection models for diploids, autotetraploids, and allotetraploids. To complement our deterministic analysis, future work should develop population genomic inference methods and stochastic models to explore the subtleties of polyploid evolution, particularly as polyploid genomic data become increasingly available. The insights from our model and similar work could have relevant applications for economically important crops, especially in light of how dominance substantially alters allele frequency dynamics and, thus, may influence selective breeding in crops with recent histories of whole genome duplication (Renny-Byfield and Wendel, 2014). As our theoretical understanding advances by incorporating additional biological complexities, these foundational insights may prove essential for predicting polyploid evolutionary trajectories in natural populations and agricultural systems.

## 5 Methods

### 5.1 Numerical Model Analysis

Our models capture the dynamical interplay between mutation and selection at a single locus in populations of diploids, autotetraploids, and allotetraploids. One of the primary objectives in this setting is the identification of points of MSB. Mathematically, these correspond to points where the change in gamete frequencies is simultaneously zero for all gamete types in the model. So, points of MSB are fixed points or equilibria solutions of the systems of ODEs.

Our model is nonlinear in both the gamete frequencies and parameters, which makes analyzing the systems of ODEs difficult; in particular, no analytical solutions exist. However, progress toward locating fixed points and evolutionary trajectories of populations can still be made. We adopt a nonlinear dynamics framework and apply classic techniques from this field to analyze our model with a focus on qualitative features such as fixed points, their stability, and bifurcations (Strogatz, 2018).

We first introduce some additional notation before sketching the numerical approaches used in analyzing the models. Let the vectorized system of ODEs be denoted as **ġ** . This vectorized function, **ġ**, will take two vector inputs, **g** and **p**, where **g** ∈ ℝ^*n−*1^ is a vector of *n* − 1 gamete frequencies and **p** ∈ ℝ^*m*^ is a vector of *m* parameters. Note that both **ġ** and **g** have *n* − 1 elements for a model with *n* gametic types because the linear constraint that the gamete frequencies must sum to 1 is first applied to the system of ODEs. For example, in the allotetraploid model, **ġ** = [*ġ*_00_, *ġ*_01_, *ġ*_11_], **g** = [*g*_00_, *g*_01_, *g*_11_], and **p** = [*s, h*_1_, *h*_2_, *h*_3_, *µ, v*].

A fixed point, **g**^*\**^ ∈ ℝ^*n−*1^, occurs where **ġ** (**g**^*\**^, **p**^*\**^) = **0** for some specific **p**^*\**^ ∈ ℝ^*m*^. Importantly, the fixed points and their stability are equivalent for the iterative map/difference equation (2) and differential equations (4). As applied to our analysis, the approximation of iterative maps by differential equations holds for weak evolutionary forces (i.e., *s, µ, v* ≪ 1). More formal analysis of this weak evolution limit has been performed in the differential equations framework (Hadeler, 1981; Hofbauer, 1985; Bürger, 1998) and in the context of replicator dynamics from evolutionary game theory (Losert and Akin, 1983).

Because no analytical solution exists for the fixed points of **ġ**, we estimate the fixed points numerically using vpasolve in Matlab2023b. We first evaluate the system of ODEs for some set of parameters by calculating **ġ** (·, **p**^*\**^). This gives a system of nonlinear, algebraic equations in the elements of **g** which is then passed to vpasolve without an initial guess. Because the systems are relatively small with three or fewer equations and, when solving **ġ** (**g**^*\**^, **p**^*\**^) = **0**, each *ġ*_*i*_ can be written as a polynomial expression in terms of the elements of **g**, vpasolve returns all (real) fixed points **g**^*\**^ for a given **p**^*\**^.

In order to classify the linear stability of each fixed point **g**^*\**^, we linearize the system about each **g**^*\**^ by keeping only O(*ε*) terms in the Taylor expansion. This is equivalent to computing the Jacobian matrix, **J** at (**g**^*\**^, **p**^*\**^). The *ij*^*th*^ element of **J** is 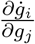 where *ġ*_*i*_ is the *i*^*th*^ differential equation of **ġ** and *g*_*j*_ is the *j*^*th*^ gamete type in **g**. The solutions of the resulting linear approximation characterized by **J** are fully specified by the eigenvalues *λ*_*i*_ and eigenvectors **v**_*i*_ of **J**. We use the Matlab function eig to numerically find the eigenvalues and eigenvectors of **J** evaluated at a fixed point.

Then, we classify the linear stability of each fixed point. In the biological context, a population near a stable gamete frequency vector will move toward **g**^*\**^ and a population near an unstable gamete frequency vector will move away from **g**^*\**^. Mathematically, a fixed point is (asymptotically) stable if the real part of all eigenvalues is negative (i.e., Re(*λ*_*i*_) *<* 0, ∀ *i* ∈ {1, …, *n* − 1}). Alternatively, a fixed point is unstable if at least one eigenvalue has positive real part (i.e., ∃ *λ*_*i*_ s.t. Re(*λ*_*i*_) *>* 0). A third, intermediate case called marginal stability can also occur when the eigenvalues all have a non-positive real part and at least one eigenvalue has an identically zero real part (i.e., Re(*λ*_*i*_) ≤ 0, ∀*i* ∈ {1, …, *n* − 1} and ∃ *λ*_*i*_ s.t. Re(*λ*_*i*_) = 0). This results in a (zero-eigenvalue) bifurcation at the fixed point.

### 5.2 Genetic Load

In our model, we use relative fitness to parameterize selection. While this is useful within each individual model, it can make explicit comparisons between populations difficult because the absolute fitness (e.g., viability or fecundity) of the most fit genotype need not be the same across populations. Instead, we calculate load, *L*, which is appropriately normalized to facilitate comparisons across models and populations. Mutation load is considered the expense, measured in terms of decreased fitness due to deleterious mutations, for the opportunity to evolve via beneficial mutations (Muller, 1950; Crow, 1970); similarly, in the case of overdominant mutations, segregation load measures the decreased fitness due to the segregation of less fit homozygous individuals in the population (Crow, 1970). We define the genetic load as the fraction by which fitness is reduced in a population of interest relative to a reference population. That is

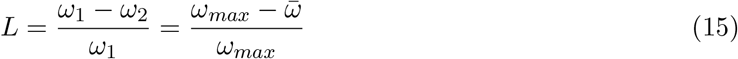

where 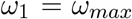 is the fitness of the reference population and *ω*_2_ = 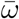 is the fitness of the population of interest. *ω*_1_ = *ω*_*max*_ is most often taken to be one which represents the fitness of a population absent any mutations (although this is not the case for overdominant mutations).

### 5.3 Discrete Simulations

To complement our numerical methods for finding fixed points or equilibria of our model, we developed discrete time simulations in Python which track the evolution of gamete frequencies over multiple generations. Concretely, our simulation follows the discrete steps of random mating, selection, meiosis, and mutation as outlined in Eqn. 1 for a set of constant parameters and a specified number of generations; however, this model assumes that the proportion of each gamete type that are produced in generation *t* is exactly equal to the proportion of each gamete type that contributes to forming the genotypes in generation *t* + 1. In mathematical terms, the genotypes for generation *t* + 1 are formed through a 1-dimensional (for autotetraploids and diploids) or 2-dimensional discrete linear convolution of the matrix of gamete frequencies formed during meiosis in generation *t*. Because this deterministic modeling approach equates the proportion of gametes formed with the proportion of gametes that contribute to the next generation, this differs from a stochastic modeling approach (used for modeling genetic drift), which would use a multinomial sampling scheme of the multiple gamete types to form the next generation. In our simulations, at the end of each generation, we record the gamete frequencies which can then be used to calculate summary statistics such as allele or genotype frequency, load, and average fitness. Scripts for the diploid, autotetraploid, and allotetraploid models and example usage is available at https://github.com/conJUSTover/Mutation Selection Balance Autos Allos.git.

## 6 Acknowledgements

This work was supported by a National Science Foundation Postdoctoral Research Fellowship in Biology (IOS-2209085 to J.L.C.) and by the National Institute of General Medical Sciences (R35 GM149235 to R.N.G.).

## Supporting Information

### S1 Detailed Supplemental Models

#### S1.1 Diploid One-Locus Model

##### S1.1.1 Notation and Random Mating

Let **g** = [*g*_0_, *g*_1_] be a two element vector of gamete frequencies, *g*_*i*_, where *i* ∈ {0, 1} is the number of derived alleles in the haploid gamete. Let **G** be a three element vector of genotype frequencies, *G*_*i*_. Under random mating, **G** is a discrete linear convolution of **g**, which we notate as **G** = **g**\***g**. Each *G*_*i*_ for *i* ∈ {0, 1, 2} is the frequency of the genotype with *i* derived alleles. Then, **G** can be written as:

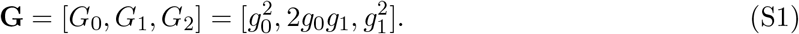

The expanded genotypes in the second vector are listed in Column 3 of Table S1 and are equivalent to the expectation of two independent draws from a binomial sample of the gametes in **g**.

**Table S1:**
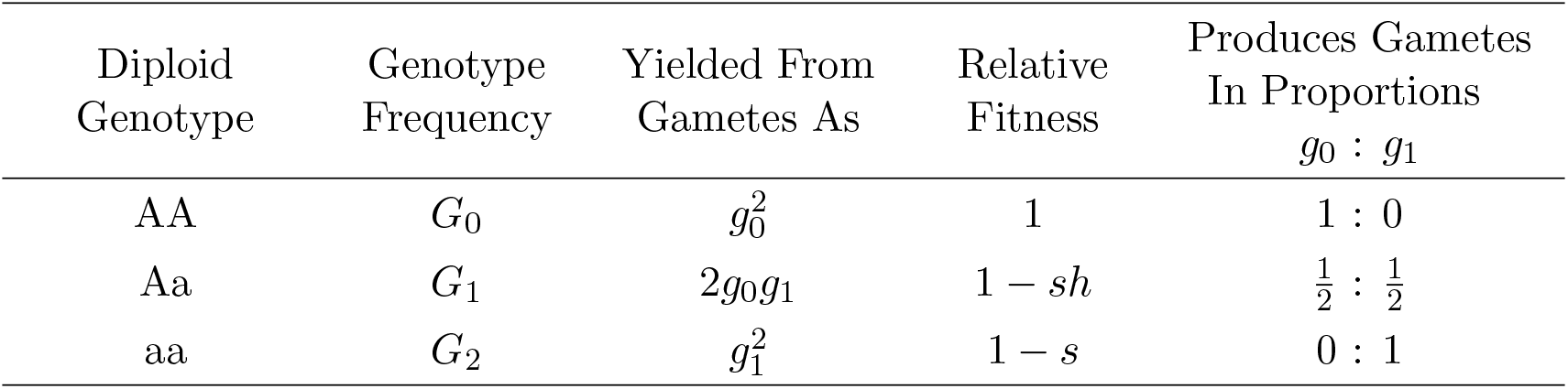
Diploid Model Summary for Mutation-Selection Balance.

##### S1.1.2 Diploid Selection

We will model the selection acting on the diploid individual with a 3-element vector of relative fitnesses, **S**, such that each element corresponds to the relative fitness acting on the corresponding element in **G** (Column 4 of Table S1). The selection coefficient is *s* and dominance for the heterozygote is denoted by *h*. Thus, the full vector of fitnesses is

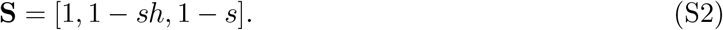

Let 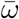 be the average fitness of the population which is the sum of the product of the *i*^*th*^ genotype frequency and *i*^*th*^ relative fitness:

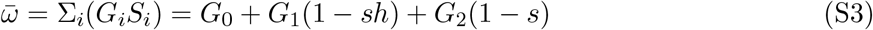

which can be simplified to

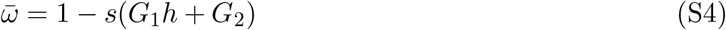

Let 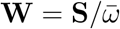 be a vector containing the normalized fitnesses, where 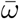 is used as the normalizing factor. Then, **W** can be written as

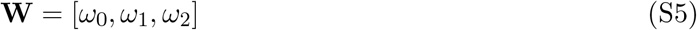

where *ω*_*i*_ denotes the normalized fitness effect acting on a genotype with *i* derived alleles.

Finally, **G**^*′*^ can be written as the Hadamard product of the genotype frequencies, **G**, and the normalized fitness effects, **W**.

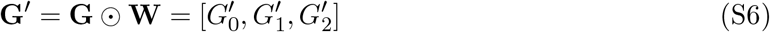

where the *i*^*th*^ element of **G**^*′*^ is simply:

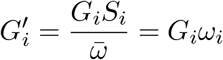

##### S1.1.3 Diploid Meiosis

We model gamete segregation with the following equations

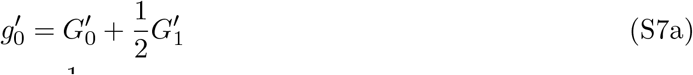

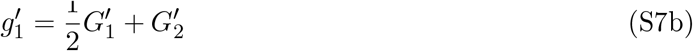

where the coefficients for each term are listed in Column 5 of Table S1.

##### S1.1.4 Diploid Mutation

The following equations model the transition probabilities between types of gametes in terms of *µ* and *v*.

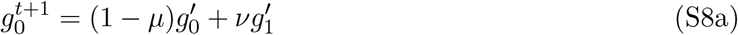

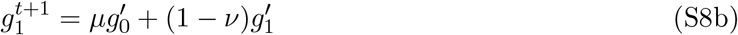

##### S1.1.5 Diploid Changes in Gamete Frequencies

To model the change in gamete frequencies between subsequent generations, we subtract a 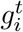 term from (S8) to obtain the system of ODEs:

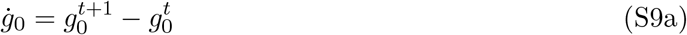

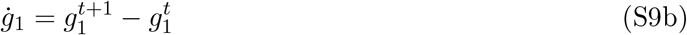

After applying the linear constraint that the sum of all gamete frequencies must equal one (i.e., Σ_*i*_*g*_*i*_ = 1), the diploid model is, by construction, equivalent to the following expanded equation which was studied by Bürger (1998) (our notation).

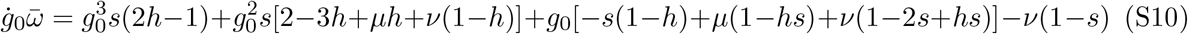

**Table S2:** Autotetraploid Model Summary for Mutation-Selection Balance.

| Autotetraploid<br>Genotype | Genotype<br>Frequency | Yielded From<br>Gametes As | Relative<br>Fitness | Produces Gametes<br>In Proportions<br>$g_0 : g_1 : g_2$ |
| --- | --- | --- | --- | --- |
| AAAA | $G_0$ | $g_0^2$ | 1 | 1 : 0 : 0 |
| AAAa | $G_1$ | $2g_0g_1$ | $1 - sh_1$ | $\frac{1}{2} : \frac{1}{2} : 0$ |
| AAaa | $G_2$ | $g_1^2 + 2g_0g_2$ | $1 - sh_2$ | $\frac{1}{6} : \frac{2}{3} : \frac{1}{6}$ |
| Aaaa | $G_3$ | $2g_1g_2$ | $1 - sh_3$ | $0 : \frac{1}{2} : \frac{1}{2}$ |
| aaaa | $G_4$ | $g_2^2$ | $1 - s$ | 0 : 0 : 1 |

#### S1.2 Autotetraploid One-Locus Model

##### S1.2.1 Notation and Random Mating

Let **g** = [*g*_0_, *g*_1_, *g*_2_] be a three element vector of gamete frequencies, *g*_*i*_, where *i* ∈ {0, 1, 2} is the number of derived alleles in each diploid gamete. Let **G** be a five element vector of genotype frequencies, *G*_*i*_. Under random mating, **G** is a discrete linear convolution of **g**, which we denote as **G** = **g**\***g**. Thus, *G*_*i*_ is the frequency of a genotype with *i* derived alleles, where *i* ∈ {0, 1, 2, 3, 4}.

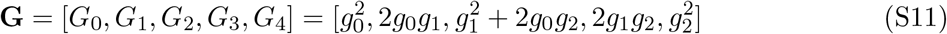

The expanded genotypes in the second vector are listed in Column 3 of Table S2 and are equivalent to the expectation of two independent draws from a multinomial sample of the gametes in **g**.

##### S1.2.2 Autotetraploid Selection

We will model selection acting on the autotetraploid individual with a vector of relative fitnesses, **S**, such that each element corresponds to the relative fitness of the corresponding element in **G** (Column 4 of Table S2. So,

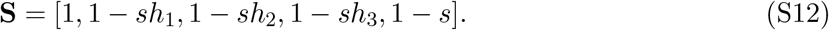

where *s* is the selection coefficient of a deleterious mutation and *h*_1_, *h*_2_, *h*_3_ are the dominance coefficients for heterozygotes with one, two, and three copies of the selected allele, respectively.

We define average fitness 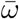 as

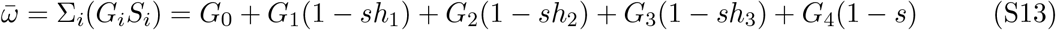

which can be simplified to

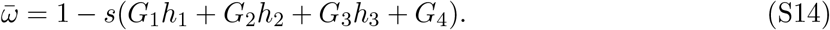

Then, the vector of normalized fitnesses is defined as 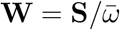 which is

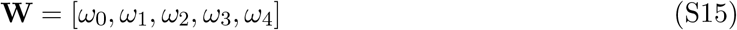

where *ω*_*i*_ is the normalized fitness effect acting on a genotype with *i* derived alleles.

Finally, the normalized genotype frequencies after selection, **G**^*′*^, are

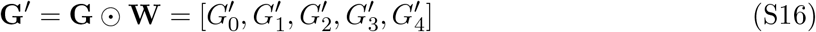

where the *i*^*th*^ element of **G**^*′*^ is simply

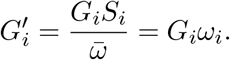

##### S1.2.3 Autotetraploid Meiosis

To model the segregation patterns of a polysomic autotetraploid (i.e., an autotetraploid with no double reduction and no preferential chromosomal pairing), we use the following system of equations

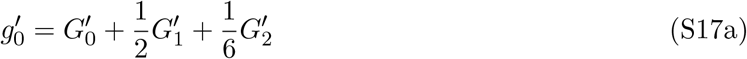

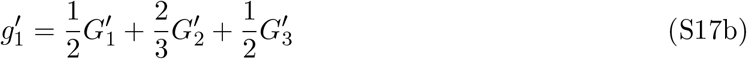

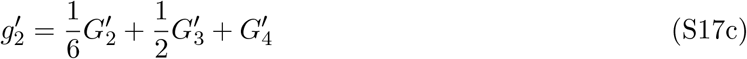

where the coefficients for each term are the gamete frequencies generated from each tetraploid genotype. These segregation frequencies are also listed in Column 5 of Table S2 and were originally derived by Haldane (1930).

##### S1.2.4 Autotetraploid Mutation

The following equations model the transition probabilities between types of gametes in terms of *µ* and *v*.

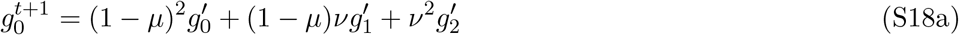

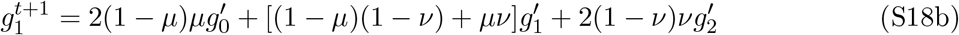

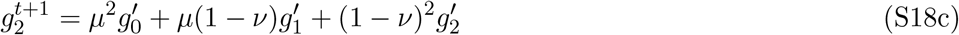

##### S1.2.5 Autotetraploid Changes in Gamete Frequencies

To model the change in gamete frequencies between subsequent generations, we subtract a 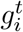 term from (S18) to obtain the following system of ODEs for autotetraploids:

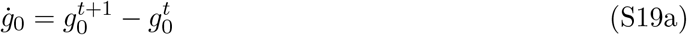

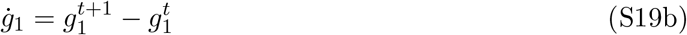

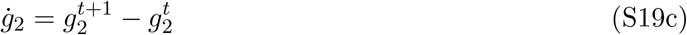

### S2 Supplement

#### S2.1 Neutral Dynamics and Panmictic Disequilibrium

Here, we consider the dynamics of each model absent mutation and selection (*s* = *µ* = *v* = 0) to isolate the approach toward Hardy-Weinberg Equilibrium (HWE). While a single generation of random mating in diploids (or within diploid subgenomes of an allopolyploid) produces genotype frequencies in HWE, polyploid populations approach HWE asymptotically, although the rate of approach differs for polyploids with disomic and tetrasomic inheritance patterns. The rate of approach can be described using panmictic disequilibrium (PD) (Gallais, 2003), a set of statistics denoted by Δ_*ij,t*_ which measure deviations from expected gamete frequencies under HWE. In general, panmictic disequilibrium is

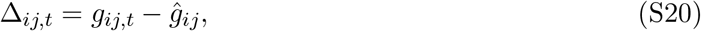

where *g*_*ij,t*_ denotes the ordered gamete frequency of type *ij* at generation *t* and *ĝ*_*ij*_ denotes the expected gamete frequency of type *ij* under HWE (Gallais, 2003). At HWE, *ĝ*_*ij*_ = *p*_*i*_*p*_*j*_ where *p*_*i*_, *p*_*j*_ are the allele frequencies of types *i, j*, respectively. Under our assumption of a biallelic locus, *p*_*i*_ = 1 − *p*_*j*_ if *i* ≠ *j*.

Δ_*ij,t*_ is closely related to linkage disequilibrium (LD) between two loci in diploids. The principal distinction is that Δ_*ij,t*_ is calculated for a single locus across multiple chromosomes, whereas LD is calculated for multiple loci on a single chromosome. Similar to how LD decays over subsequent generations and recombination events, so too does Δ_*ij,t*_ decay over time. However, this decay is dependent on the proportion of gametes which are identical by descent (IBD) from the previous generation instead of the recombination rate between loci (Slatkin, 2008). Additionally, for biallelic loci, just as two-locus gamete frequencies can be written in terms of allele frequencies and a single LD statistic, we can recover gamete frequencies from allele frequencies and a single PD statistic. Concretely, Δ_*ii,t*_ = Δ_*jj,t*_ = −Δ_*ij,t*_ = −Δ_*ji,t*_ for allelic types *i* ≠ *j*.

Let *c* denote the expected proportion of an individual’s gametes which reproduce the allelic combination in one of the gametes that formed that individual (i.e., *c* is the proportion of gametes which are IBD). Then, 1 − *c* of the gametes are not IBD. So,

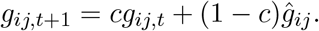

Subtracting a *ĝ*_*ij*_ term from both sides gives

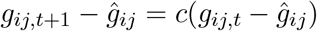

which, substituting (S20) is equivalent to

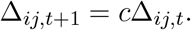

Thus, it is clear that Δ_*ij*_ decays geometrically at rate *c* (Gallais, 2003). For a panmictic tetraploid population with polysomic inheritance, this set of statistics has been shown to decay geometrically with a rate of *c* = 1*/*3 (Haldane, 1930) because two in six (or one in three) gametes are IBD in an autotetraploid individual.

For a biallelic locus in an allotetraploid individual, there are only four possible gametes (compared to six for autotetraploids) because chromosomes from the same subgenome cannot segregate to the same gamete. Two of these four gametes are IBD to the gametes which formed the allote-traploid individual; thus, the proportion of gametes which are IBD is one in two, and Δ_*ij,t*_ decays geometrically at a rate of *c* = 1*/*2. We emphasize that while the marginal genotypes of the (diploid) subgenomes each reach HWE in a single generation (as in standard diploid populations), this geometric decay is associated with the asymptotic breakdown of the correlation of genotype frequencies between subgenomes and, thus, applies only to the overall genotypes of the allotetraploid (Gerard, 2023*a*). Here, we see that an autotetraploid population will approach HWE 33% faster than an allotetraploid population, but any substantive amount of panmictic disequilibrium is removed by random mating in only a few generations (Fig. S1).

**Figure S1:**
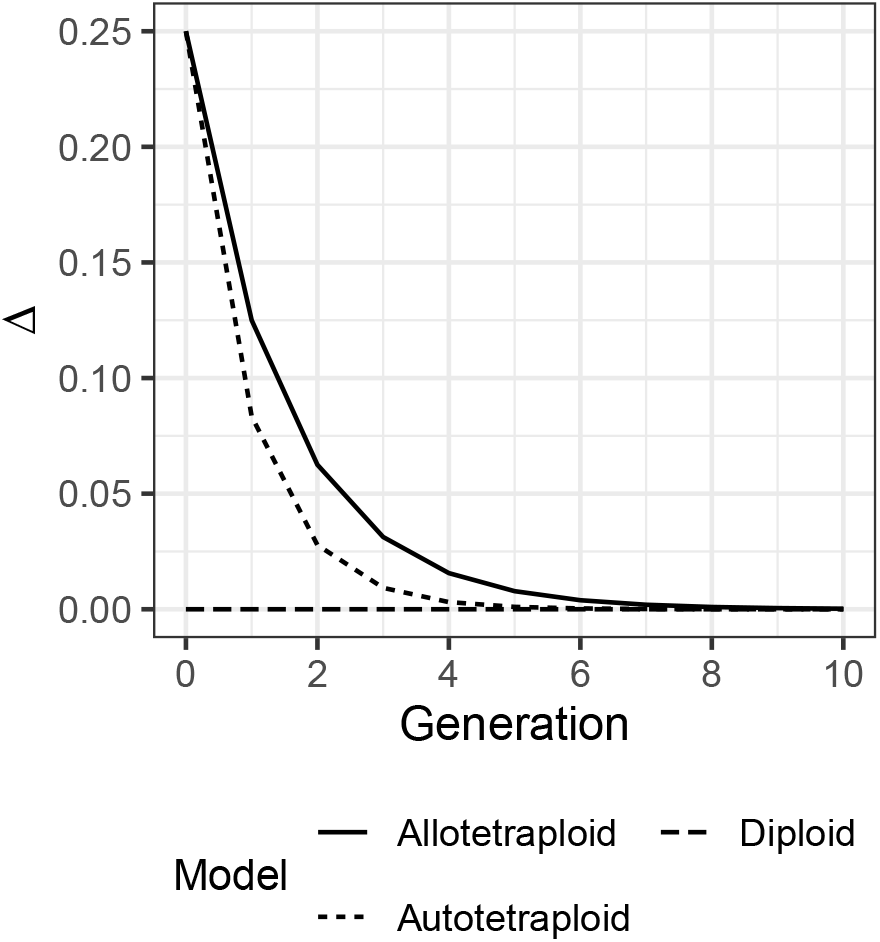
Decay of panmictic disequilibrium over time. We plot the panmictic disequilbrium for homozygous gametes (Δ_00_ = Δ_11_). Populations are simulated with neither selection nor mutation and start with *p* = *q* = 0.5 and only homozygous gametes present in the first generation.

#### S2.2 Figures

**Figure S2:**
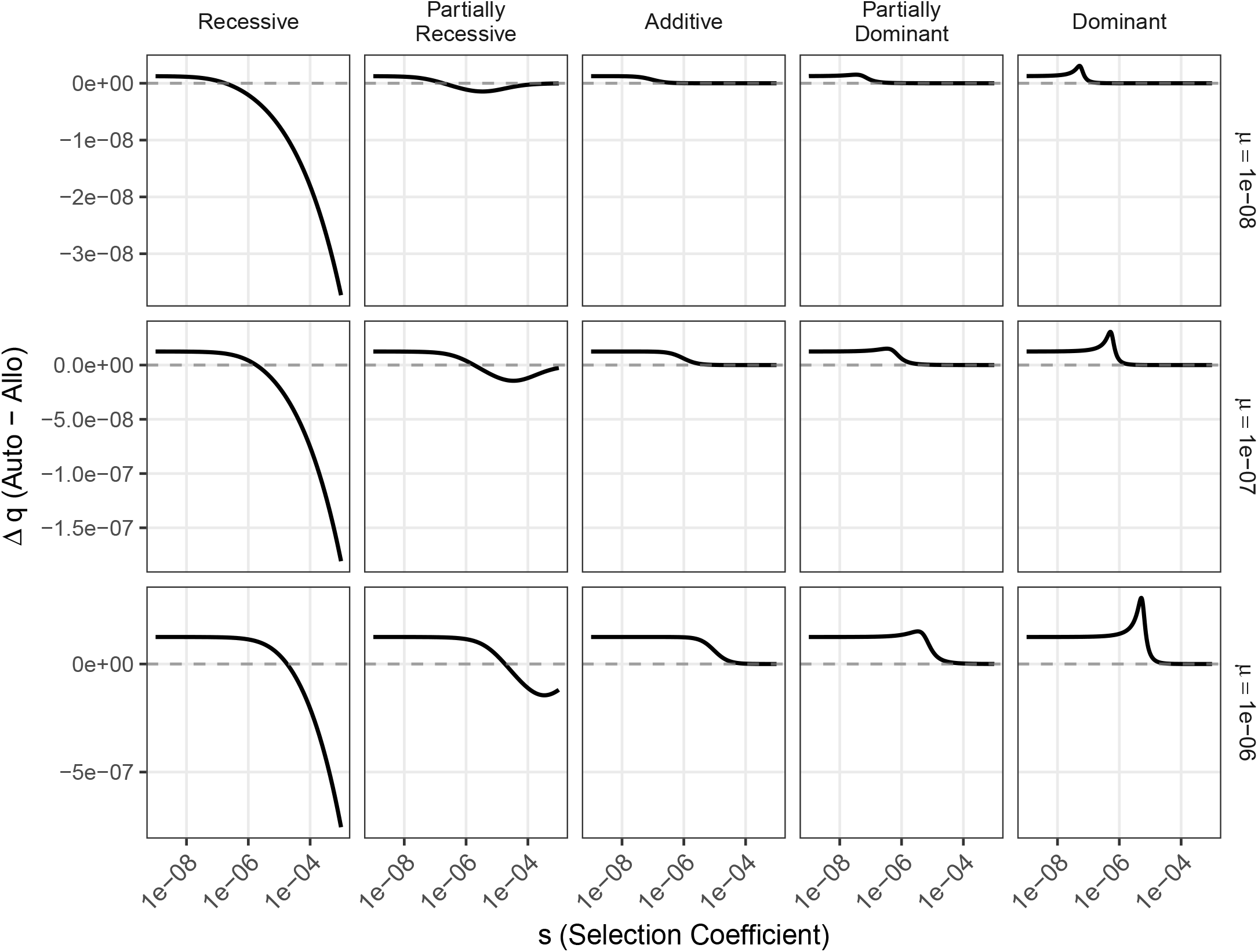
Differences between auto- and allotetraploid models at mutation-selection equilibria across dominance relations and mutation rates. Differences in allele frequency are shown. For all panels, *µ* = *ν*. The difference in equilibrium allele frequency between the tetraploid models is of order *µ* (i.e., Δ*q* ≈ *O*(*µ*)). Dominance coefficients for the partially recessive column are *h*_1_ = 0.02, *h* = *h*_2_ = 0.1, and *h*_3_ = 0.4 and for the partially dominant column are *h*_1_ = 0.6, *h* = *h*_2_ = 0.9, and *h*_3_ = 0.98.

**Figure S3:**
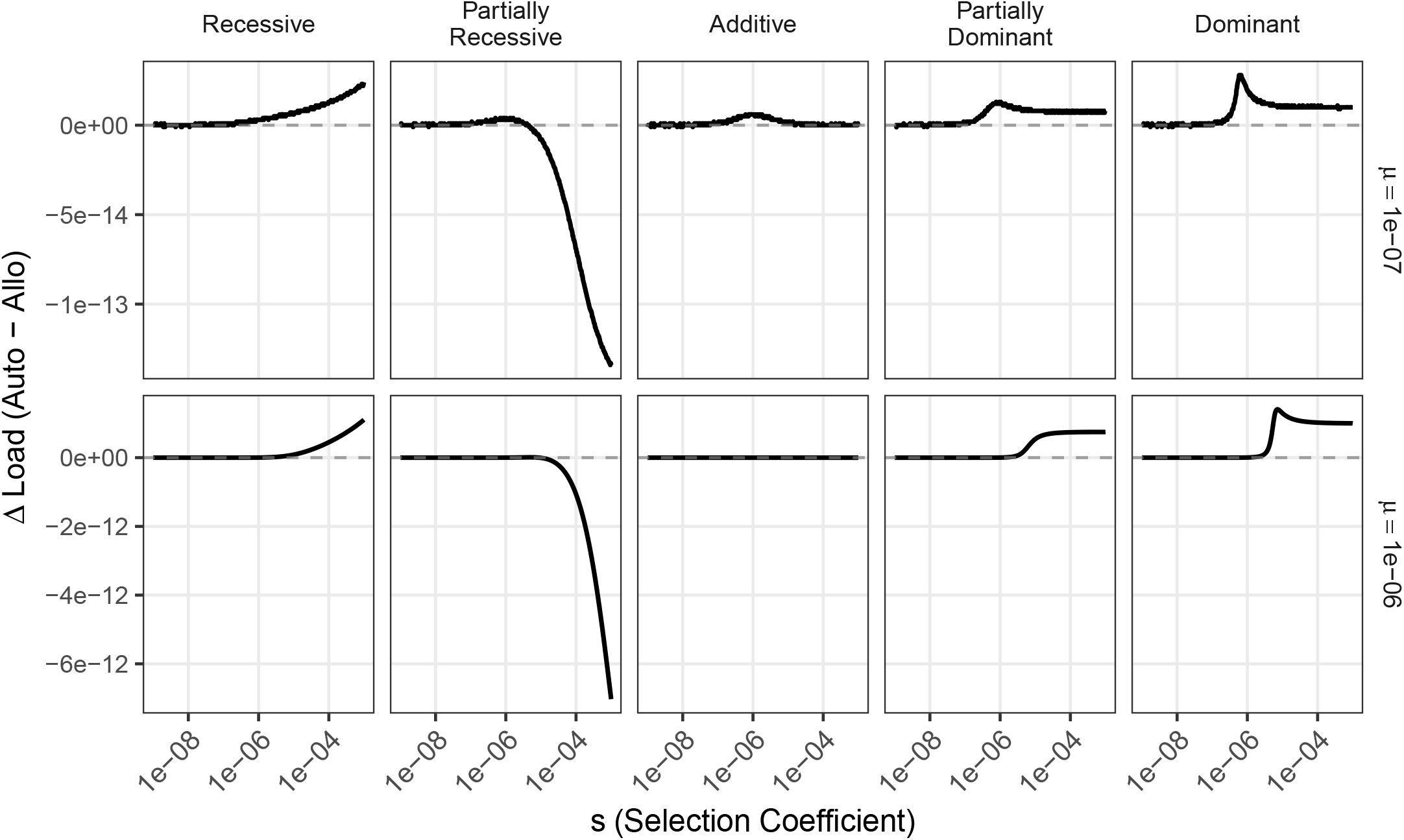
Differences between auto- and allotetraploid models at mutation-selection equilibria across dominance relations and mutation rates. Differences in mutation load are shown. For all panels, *µ* = *v*. The *µ* = 10^*−*8^ is not shown because the differences in load fall within the numerical precision of doubles. Dominance coefficients for the partially recessive column are *h*_1_ = 0.02, *h* = *h*_2_ = 0.1, and *h*_3_ = 0.4 and for the partially dominant column are *h*_1_ = 0.6, *h* = *h*_2_ = 0.9, and *h*_3_ = 0.98.

**Figure S4:**
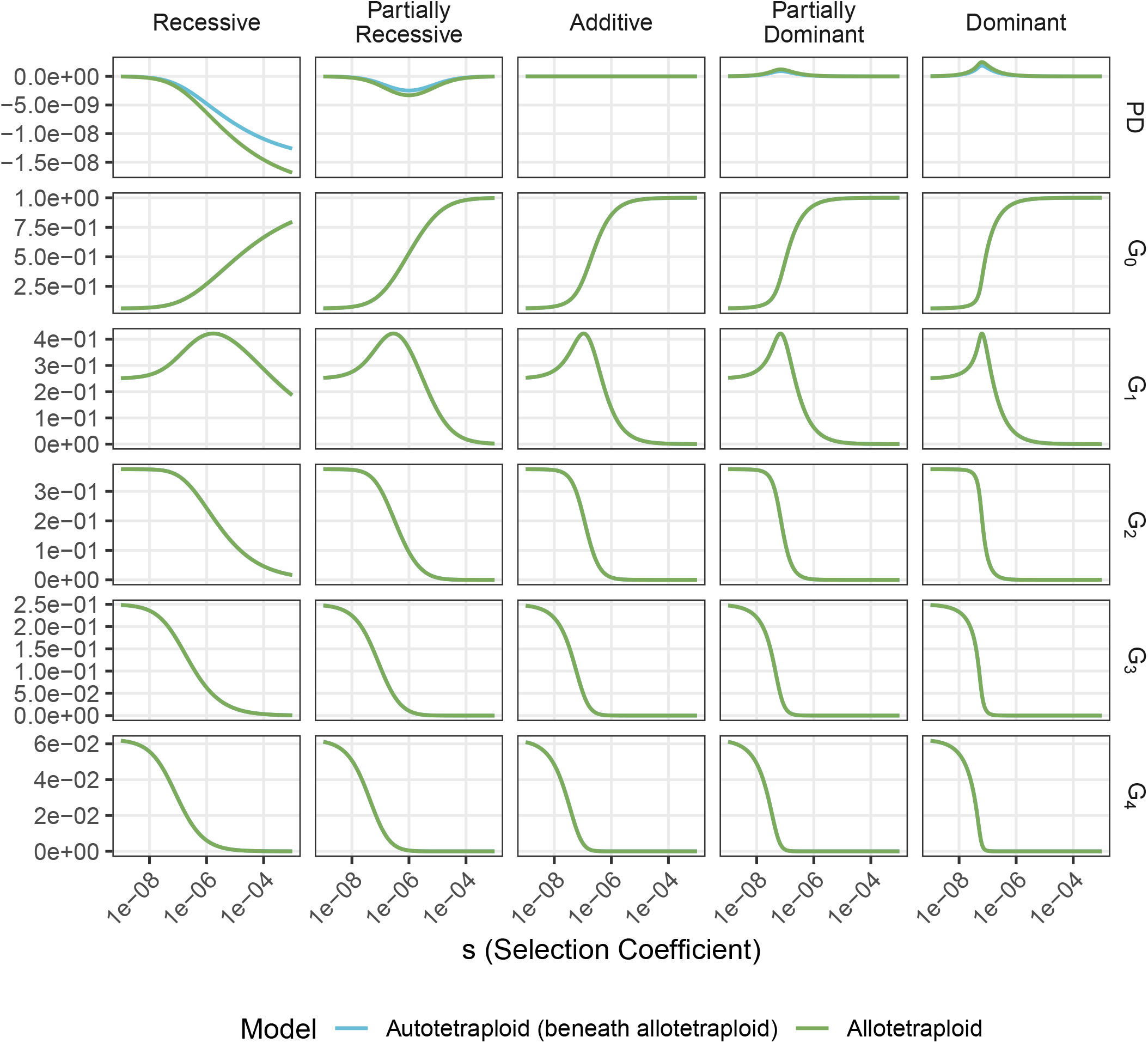
Panmictic disequilibrium (PD) and genotype frequencies at mutation-selection balance for the same parameters as Fig. 1. Genotypes for allotetraploids are collapsed by summing over the two indices for each subgenome (e.g., *G*_1_ ≡ *G*_01_ + *G*_10_). Of particular note are the local maxima of *G*_1_ which, for fully dominant mutations, have an outsized impact on the mutation load of the population, leading to the bump and non-monotonic behavior in Fig. 1J near *s* = 10^*−*7^. This bump and non-monotonic behavior are not present in the diploid model because the genotypes are never distributed such that there are more heterozygotes with a single derived allele than homozygotes with no derived alleles for *q* ≤ .5. Here, the presence of PD can further distort the genotype frequencies and also contributes to the mutation load.

**Figure S5:**
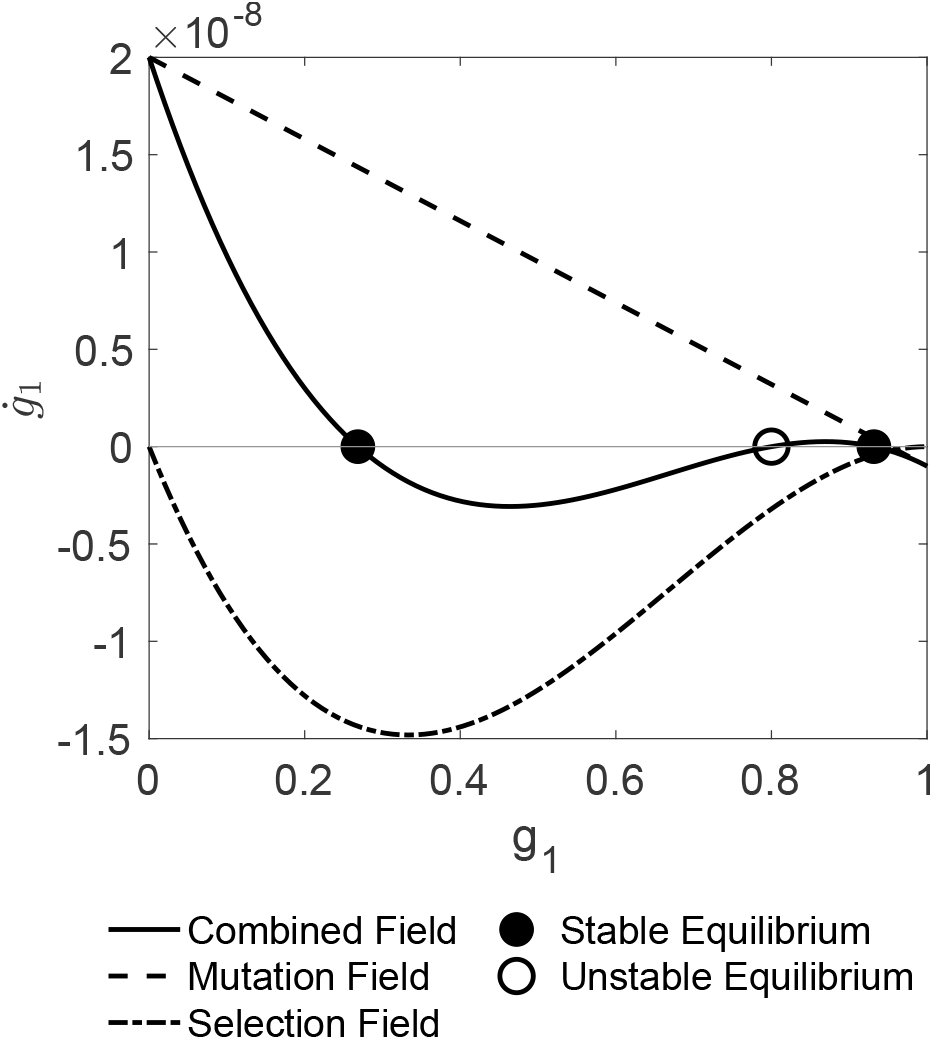
Phase space for diploids with *s* = 1*e*^*−*7^, *h* = 1, *µ* = 2*e*^*−*8^, and *ν* = 1*e*^*−*9^. Equilibria occur where the magnitude of the mutation and selection fields are equal but opposite in sign.

**Figure S6:**
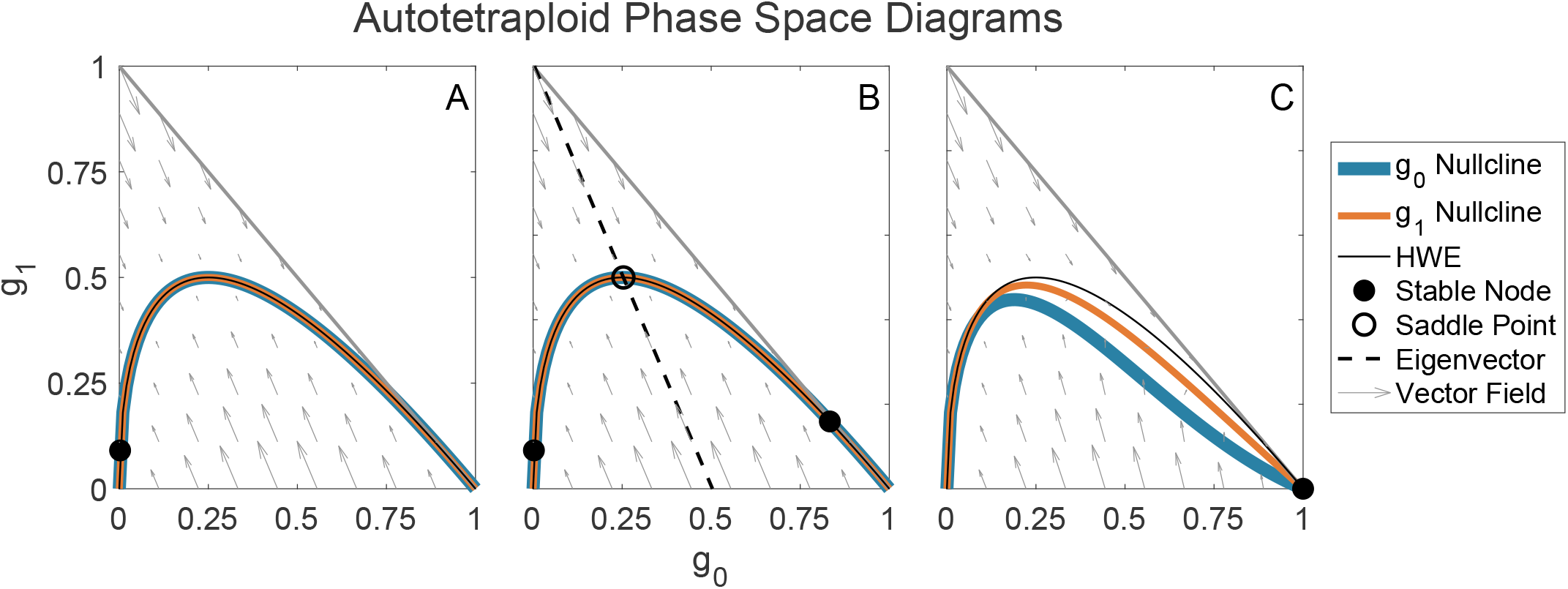
Phase space for autotetraploids with *h*_1_ = *h*_2_ = *h*_3_ = 1, *µ* = 2*e*^*−*8^, and *ν* = 1*e*^*−*9^. For A, *s* = 1*e*^*−*9^, B *s* = 3*e*^*−*7^, and C *s* = 0.3. This shows how the equilibria structure changes as parameters are varied. In between each panel, a bifurcation occurs causing the number of equilibria to change. In B, the dashed line is an eigenvector which approximately separates the phase space into two basins of attraction such that a population to the left of this line will move toward the stable equilibria near *g*_0_ = 0 and vice versa.

**Figure S7:**
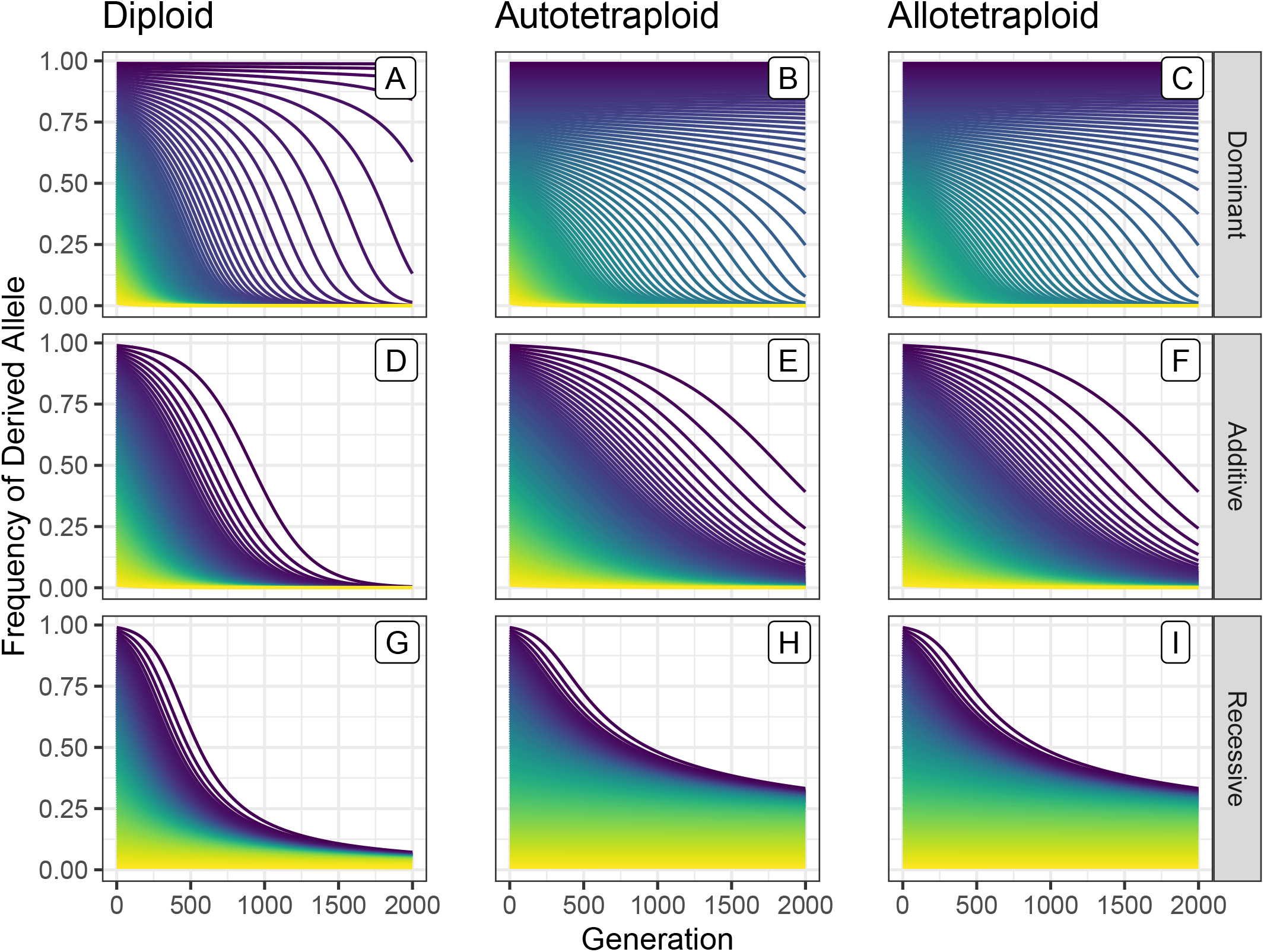
Trajectories of allele frequency over time for *s* = .01, *µ* = 1*e*^*−*8^, and *ν* = 1*e*^*−*8^. All populations begin in Hardy-Weinberg Equilibrium with an initial allele frequency from 0 to 1. The coloration denotes the initial allele frequency when the simulation started (i.e., at *t* = 0).

**Figure S8:**
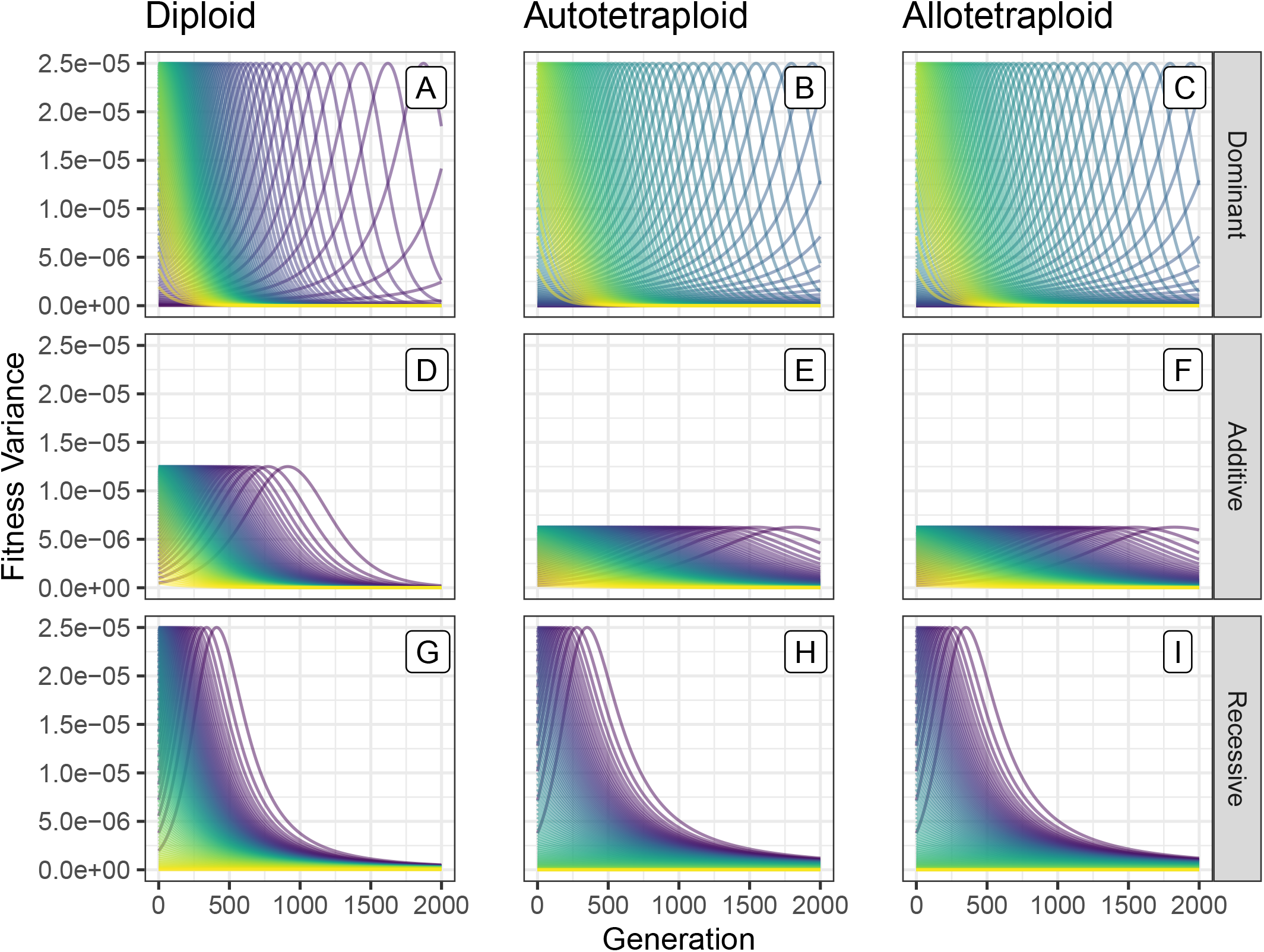
Trajectories of fitness variance over time for *s* = .01, *µ* = 1*e*^*−*8^, and *ν* = 1*e*^*−*8^. All populations begin in Hardy-Weinberg Equilibrium with an initial allele frequency from 0 to 1. The coloration denotes the initial allele frequency when the simulation started (i.e., at *t* = 0).

**Figure S9:**
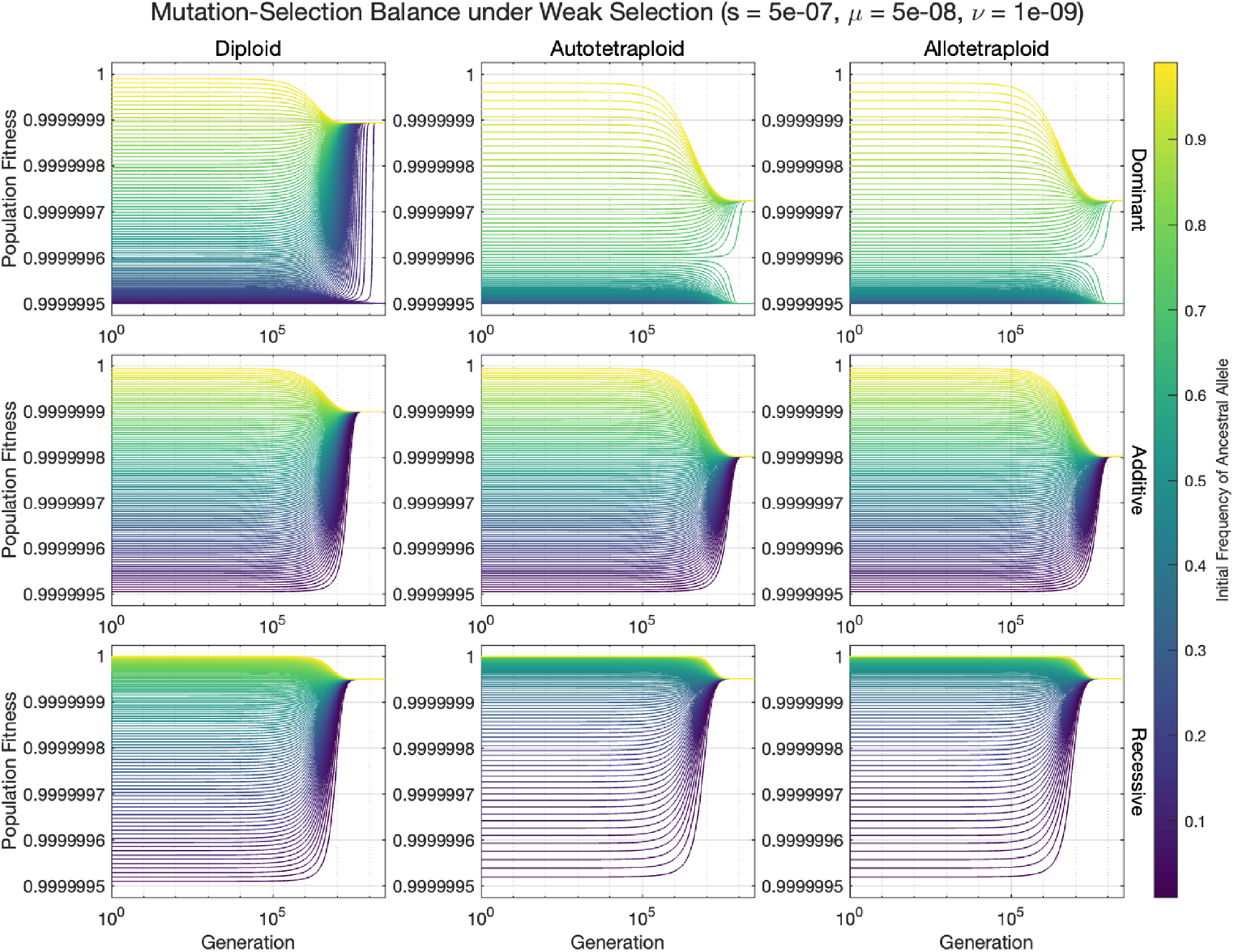
Trajectories of allele frequency over time for *s* = 5 × 10^*−*7^, *µ* = 5 × 10^*−*8^, and *v* = 10^*−*9^. All populations begin in Hardy-Weinberg Equilibrium with an initial allele frequency between 0 to 1. The coloration denotes the initial allele frequency when the simulation started (i.e., at *t* = 0). The trajectories are calculated by applying a standard stiff solver to the log-transformed system of ODEs (specifically we utilize the ode15s function in Matlab).

**Figure S10:**
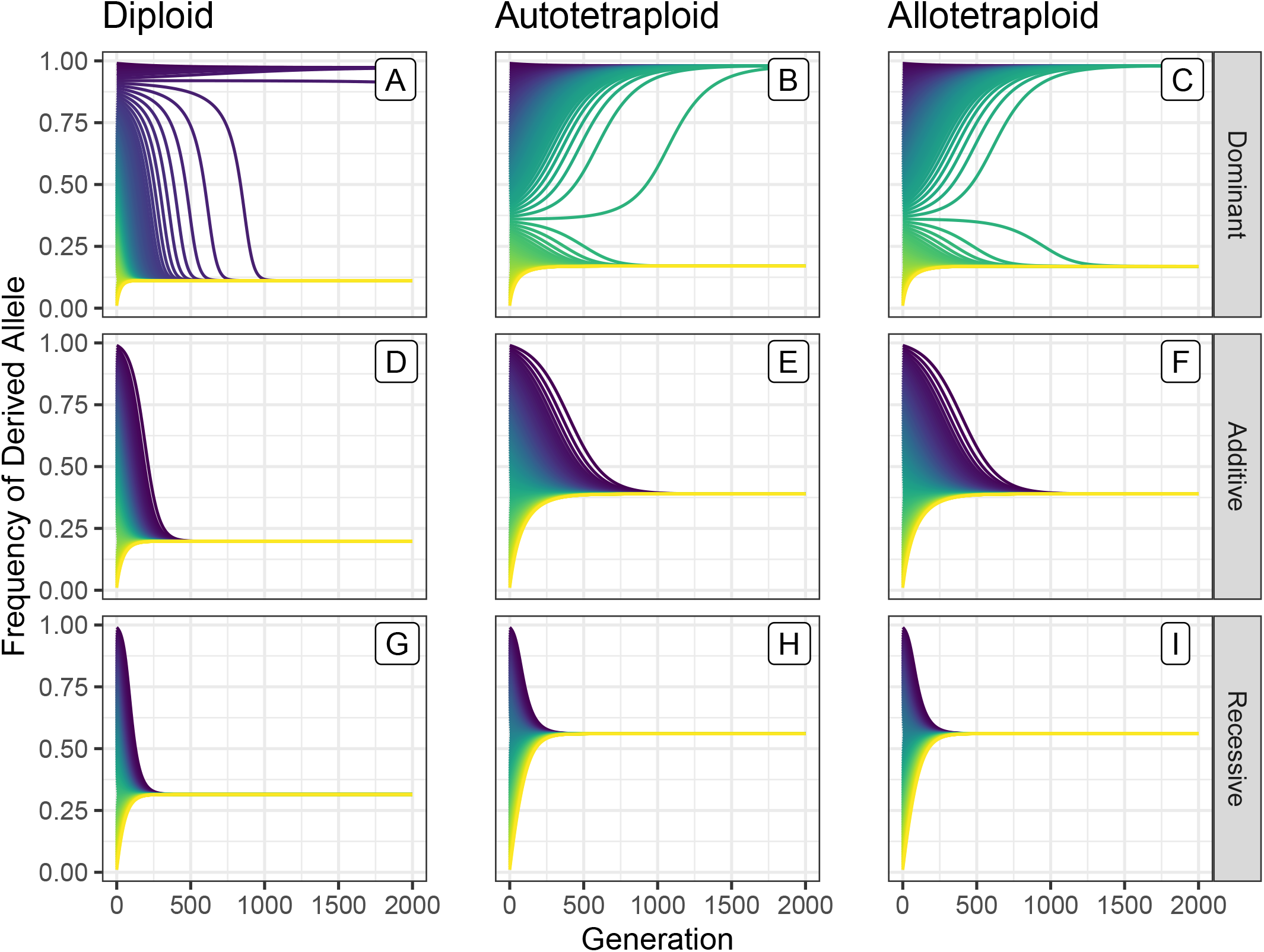
Trajectories of allele frequency over time for *s* = .05, *µ* = 5 × 10^*−*3^, and *ν* = 10^*−*4^. All populations begin in Hardy-Weinberg Equilibrium with an initial allele frequency between 0 to 1. The coloration denotes the initial allele frequency when the simulation started (i.e., at *t* = 0).

**Figure S11:**
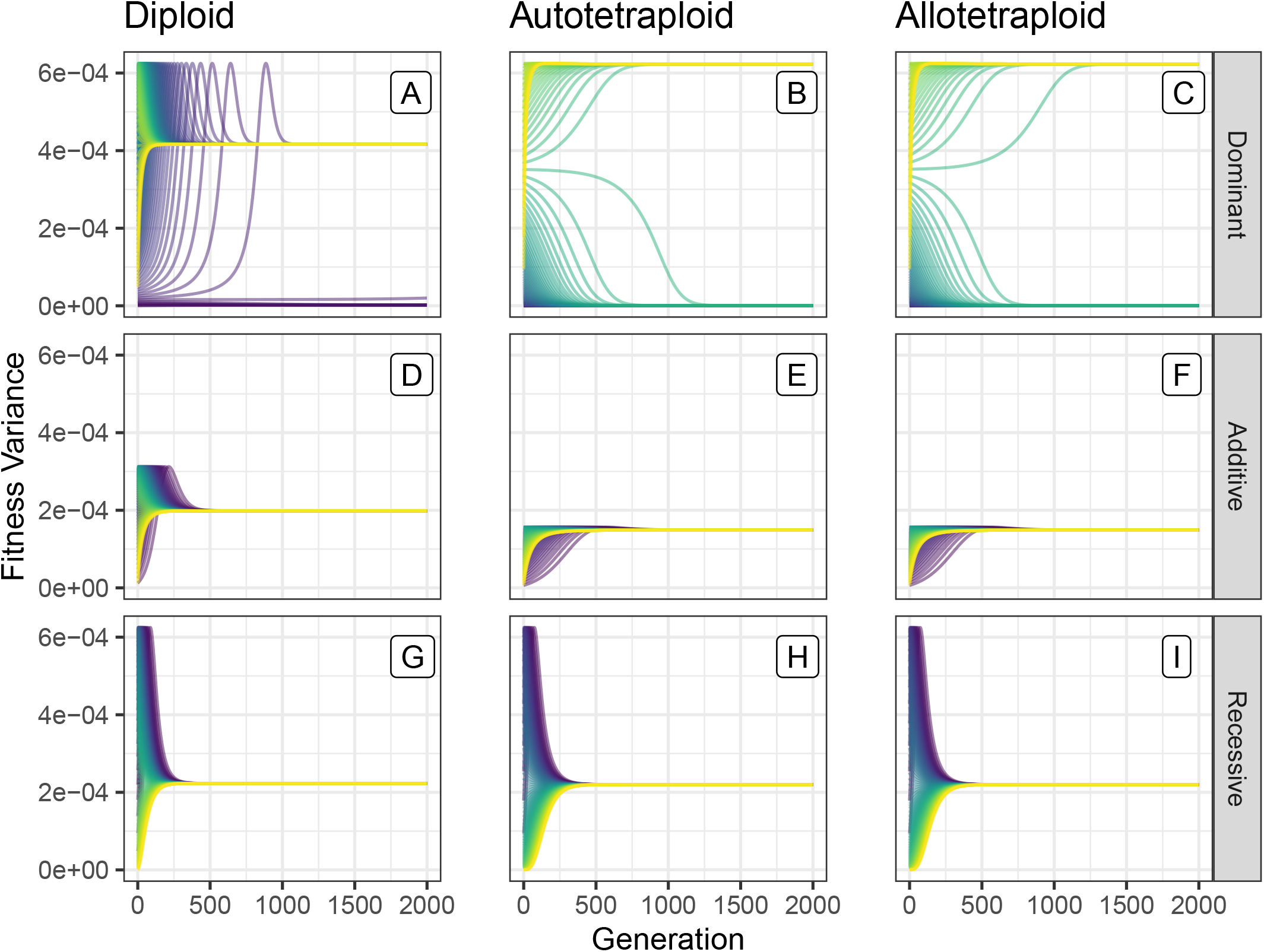
Trajectories of fitness variance over time for *s* = .05, *µ* = 5 × 10^*−*3^, and *ν* = 10^*−*4^. All populations begin in Hardy-Weinberg Equilibrium with an initial allele frequency between 0 to 1. The coloration denotes the initial allele frequency when the simulation started (i.e., at *t* = 0).

**Figure S12:**
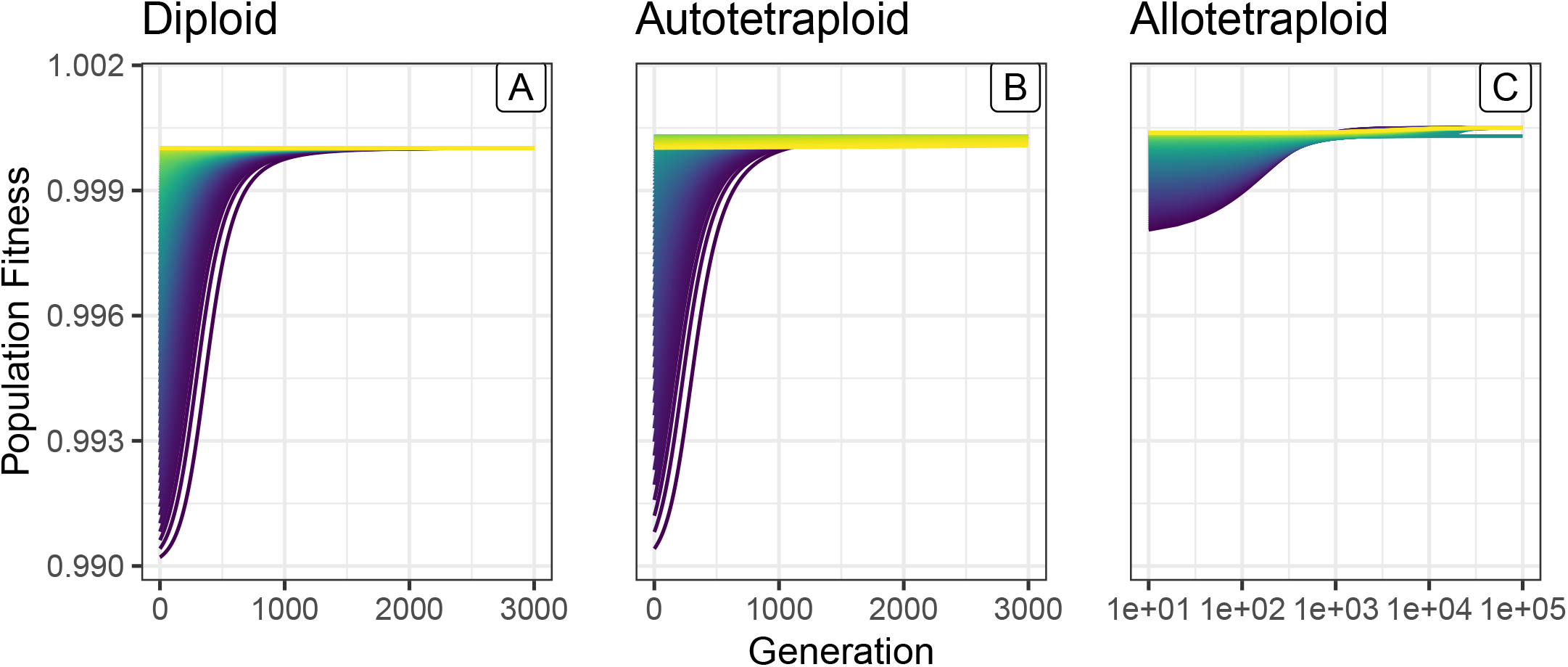
Trajectories of average fitness over time for *s* = 0.01, *h* = *h*_1_ = *h*_2_ = *h*_3_ = −0.05, and *µ* = *v* = 10^*−*8^. Each trajectory shows the evolution of a single simulation beginning in Hardy-Weinberg Equilibrium with coloration denoting the initial allele frequency when the simulation started (i.e., at *t* = 0). For the diploids and autotetraploids, initial allele frequencies were chosen in increments of 0.01 ranging from 0.01 to 0.99. For the allotetraploid simulations, initial allele frequencies in one subgenome were incremented by 0.01 ranging from 0.01 to 0.99 and were held fixed at 0.5 in the second subgenome.

**Figure S13:**
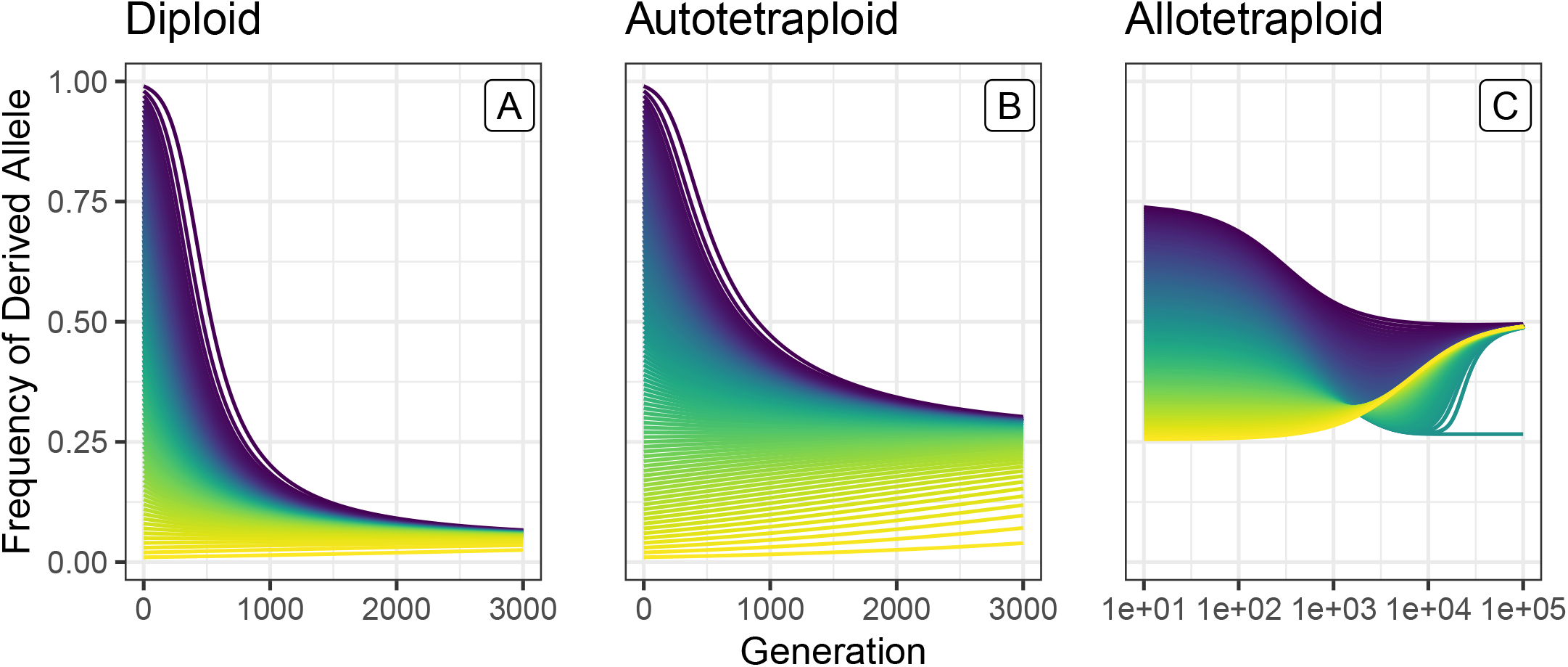
Trajectories of allele frequency over time for *s* = 0.01, *h* = *h*_1_ = *h*_2_ = *h*_3_ = −0.05, and *µ* = *v* = 10^*−*8^. Each trajectory shows the evolution of a single simulation beginning in Hardy-Weinberg Equilibrium with coloration denoting the initial allele frequency when the simulation started (i.e., at *t* = 0). For the diploids and autotetraploids, initial allele frequencies were chosen in increments of 0.01 ranging from 0.01 to 0.99. For the allotetraploid simulations, initial allele frequencies in one subgenome were incremented by 0.01 ranging from 0.01 to 0.99 and were held fixed at 0.5 in the second subgenome.

**Figure S14:**
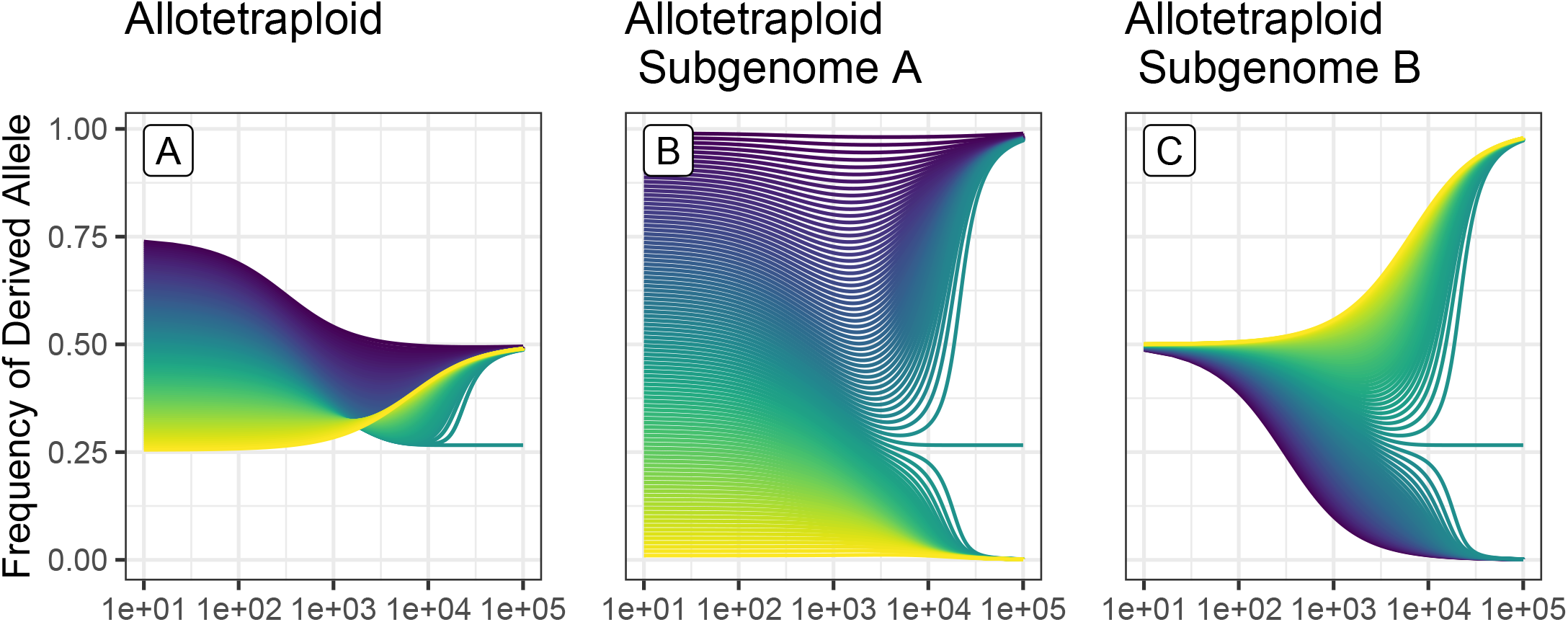
Trajectories of global and subgenome allele frequencies over time for *s* = 0.01, *h* = *h*_1_ = *h*_2_ = *h*_3_ = −0.05, and *µ* = *v* = 10^*−*8^. Each trajectory shows the evolution of a single simulation beginning in Hardy-Weinberg Equilibrium with coloration denoting the initial allele frequency when the simulation started (i.e., at *t* = 0). Initial allele frequencies in one subgenome were incremented by 0.01 ranging from 0.01 to 0.99 and were held fixed at 0.5 in the second subgenome.

